# Environment Classification for Robotic Leg Prostheses and Exoskeletons using Deep Convolutional Neural Networks

**DOI:** 10.1101/2021.06.24.449600

**Authors:** Brokoslaw Laschowski, William McNally, Alexander Wong, John McPhee

**Affiliations:** Department of Systems Design Engineering, University of Waterloo, Waterloo ON, Canada; Waterloo Artificial Intelligence Institute, University of Waterloo, Waterloo ON, Canada

**Keywords:** computer vision, deep learning, exoskeletons, rehabilitation robotics, prosthetics, wearables, artificial intelligence, biomechatronics

## Abstract

Robotic leg prostheses and exoskeletons can provide powered locomotor assistance to older adults and/or persons with physical disabilities. However, the current locomotion mode recognition systems being developed for intelligent high-level control and decision-making use mechanical, inertial, and/or neuromuscular data, which inherently have limited prediction horizons (i.e., analogous to walking blindfolded). Inspired by the human vision-locomotor control system, we designed and evaluated an advanced environment classification system that uses computer vision and deep learning to forward predict the oncoming walking environments prior to physical interaction, therein allowing for more accurate and robust locomotion mode transitions. In this study, we first reviewed the development of the ExoNet database – the largest and most diverse open-source dataset of wearable camera images of indoor and outdoor real-world walking environments, which were annotated using a hierarchical labelling architecture. We then trained and tested over a dozen state-of-the-art deep convolutional neural networks (CNNs) on the ExoNet database for large-scale image classification of the walking environments, including: EfficientNetB0, InceptionV3, MobileNet, MobileNetV2, VGG16, VGG19, Xception, ResNet50, ResNet101, ResNet152, DenseNet121, DenseNet169, and DenseNet201. Lastly, we quantitatively compared the benchmarked CNN architectures and their environment classification predictions using an operational metric called NetScore, which balances the image classification accuracy with the computational and memory storage requirements (i.e., important for onboard real-time inference). Although we designed this environment classification system to support the development of next-generation environment-adaptive locomotor control systems for robotic prostheses and exoskeletons, applications could extend to humanoids, autonomous legged robots, powered wheelchairs, and assistive devices for persons with visual impairments.

## 1 Introduction

There are currently hundreds of millions of individuals worldwide with mobility impairments resulting from aging and/or physical disabilities. For example, the number of people with limb amputations in the United States alone is expected to more than double from ~1.6 million in 2005 to ~3.6 million in 2050 (Ziegler-Graham et al., 2008); this increased prevalence is primarily driven by the aging population and higher rates of dysvascular diseases, especially diabetes, among older adults. The World Health Organization recently projected that the number of individuals above 65 years will increase from ~524 million in 2010 to ~1.5 billion in 2050 (Grimmer et al., 2019). The global prevalence of musculoskeletal disorders like osteoarthritis is ~151 million and neurological conditions like Parkinson’s disease, cerebral palsy, and spinal cord injury affect approximately 5.2 million, 16 million, and 3.5 million persons, respectively (Grimmer et al., 2019). Moreover, the number of individuals needing physical rehabilitation due to immobility and/or disease has recently been exacerbated by the coronavirus disease 2019 (COVID-19) (Negrini et al., 2021). Mobility impairments, especially those resulting in wheelchair dependency, are often associated with reduced independence and secondary health complications, including osteoporosis, coronary artery disease, obesity, hypertension, and sarcopenia (Grimmer et al., 2019).

Fitted with conventional passive assistive devices, individuals with mobility impairments tend to fall more frequently and walk more energetically inefficient and with greater biomechanical asymmetries compared to healthy young adults, which have implications on joint and bone degeneration resulting from disproportionate loading on the unaffected limb (i.e., assuming unilateral impairments) (Tucker et al., 2015). One reason for these gait abnormalities is that many locomotor activities, like climbing stairs, require net positive mechanical work about the lower-limb joints via power generation from the human muscles. However, limb amputation or paralysis removes/diminishes these sources of biological actuation, therein compromising the user’s ability to effectively and efficiently perform many locomotor activities of daily living. Fortunately, robotic leg prostheses and exoskeletons can replace the propulsive function of the amputated or impaired biological limbs and allow users to perform locomotor activities that require net positive mechanical work by using motorized hip, knee, and/or ankle joints (Krausz and Hargrove, 2019; Laschowski and Andrysek, 2018; Tucker et al., 2015; Young and Ferris, 2017; Zhang et al., 2019a). However, control of these robotic assistive devices is exceptionally difficult and often considered one of the leading challenges to real-world deployment (Tucker et al., 2015; Young and Ferris, 2017).

Most robotic prosthesis and exoskeleton control systems for human locomotion involve a hierarchical architecture, including high, mid, and low-level controllers (Tucker et al., 2015; Young and Ferris, 2017) (see Figure 1). The high-level controller is responsible for state estimation and inferring the user’s locomotor intent (e.g., climbing stairs, sitting down, or level-ground walking). The mid-level controller converts the locomotor activity into mode-specific reference trajectories often using dynamic equations of the human-robot system; this level of control consists of individual finite-state machines with discrete mechanical impedance parameters like stiffness and damping coefficients, which are heuristically tuned for different locomotor activities to generate the desired actuator joint torques (i.e., the desired device state). Tunning these system parameters can be time-consuming as the number of parameters increases with the number of states per locomotor activity, the number of activity modes, and the number of actuated joints. The low-level controller uses standard controls engineering algorithms like proportional-integral-derivative (PID) control to calculate the error between the measured and desired device states and command the robotic actuators to minimize the error using reference tracking and closed-loop feedback control (Krausz and Hargrove, 2019; Tucker et al., 2015; Young and Ferris, 2017; Zhang et al., 2019a).

**Figure 1.**
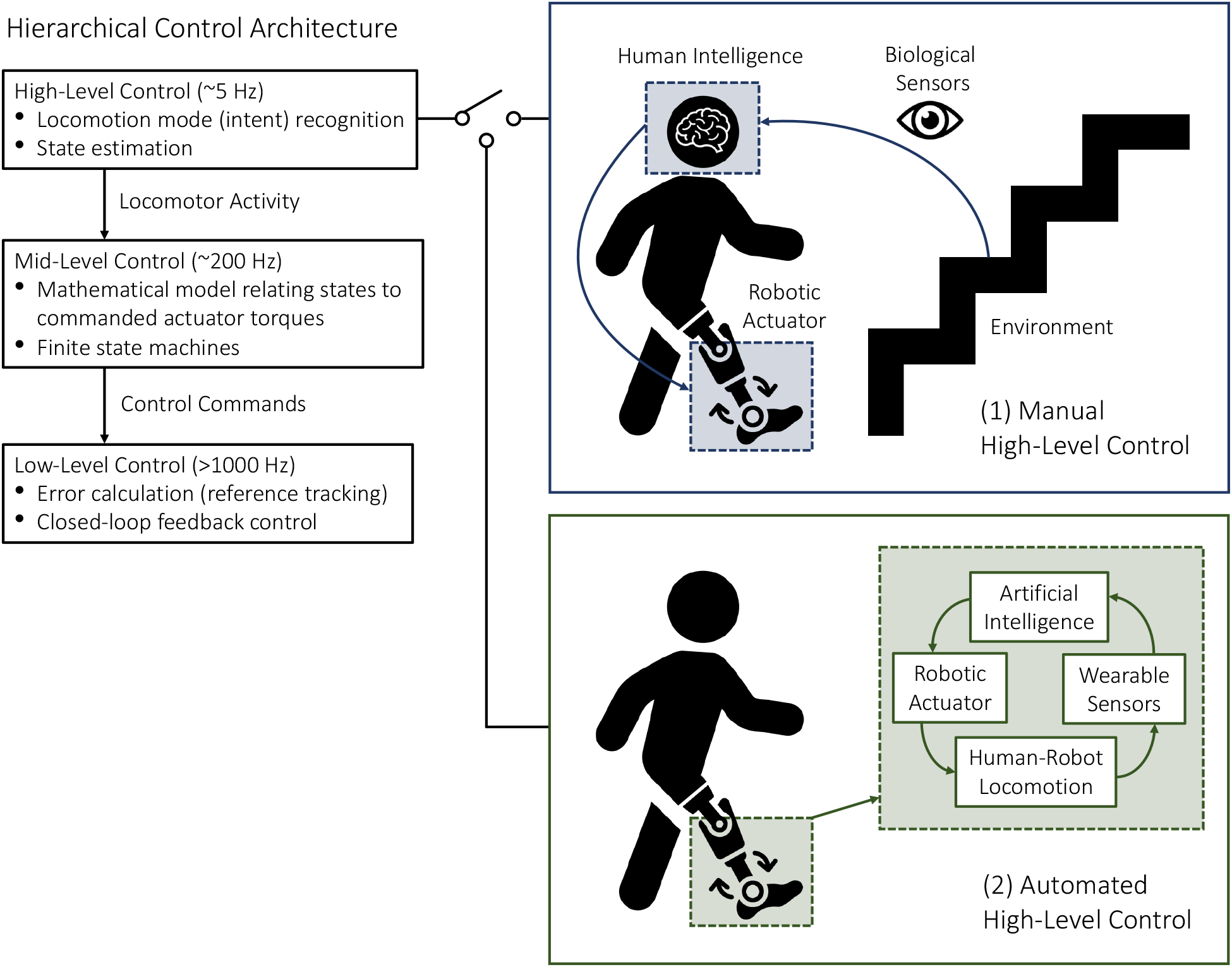
Hierarchical control architecture of robotic leg prostheses and exoskeletons, including high, mid, and low-level controllers. The high-level controller selects the desired locomotion mode using either (1) manual communication from the user (i.e., for commercially available devices) or (2) automated systems (i.e., for devices under research and development).

Nonrhythmic movements and high-level transitions between different locomotor activities remain significant challenges. Inaccurate and/or delayed decisions could result in loss-of-balance and injury, which can be especially problematic when involving stairs. Switching between different mid-level controllers is supervised by the high-level controller, which infers the locomotor intent using either sensor data (i.e., for devices under research and development) or direct communication from the user (i.e., for commercially available devices) (see Figure 1). Commercial devices like the Össur Power Knee prosthesis and the ReWalk and Indego powered lower-limb exoskeletons require the users to perform exaggerated movements or use hand controls to manually switch between locomotion modes (Tucker et al., 2015; Young and Ferris, 2017). Although highly accurate in communicating the user’s locomotor intent, such manual high-level control and decision making can be time-consuming, inconvenient, and cognitively demanding (Karacan et al., 2020). Researchers have thus been working on developing automated locomotion mode recognition systems using pattern recognition algorithms and data from wearable sensors like inertial measurement units (IMUs) and surface electromyography (EMG), therein shifting the high-level control burden from the user to an intelligent controller (Tucker et al., 2015; Young and Ferris, 2017) (see Figure 2). The ideal automated high-level controller would estimate the individual states and interactions within the overall human-robot-environment system.

**Figure 2.**
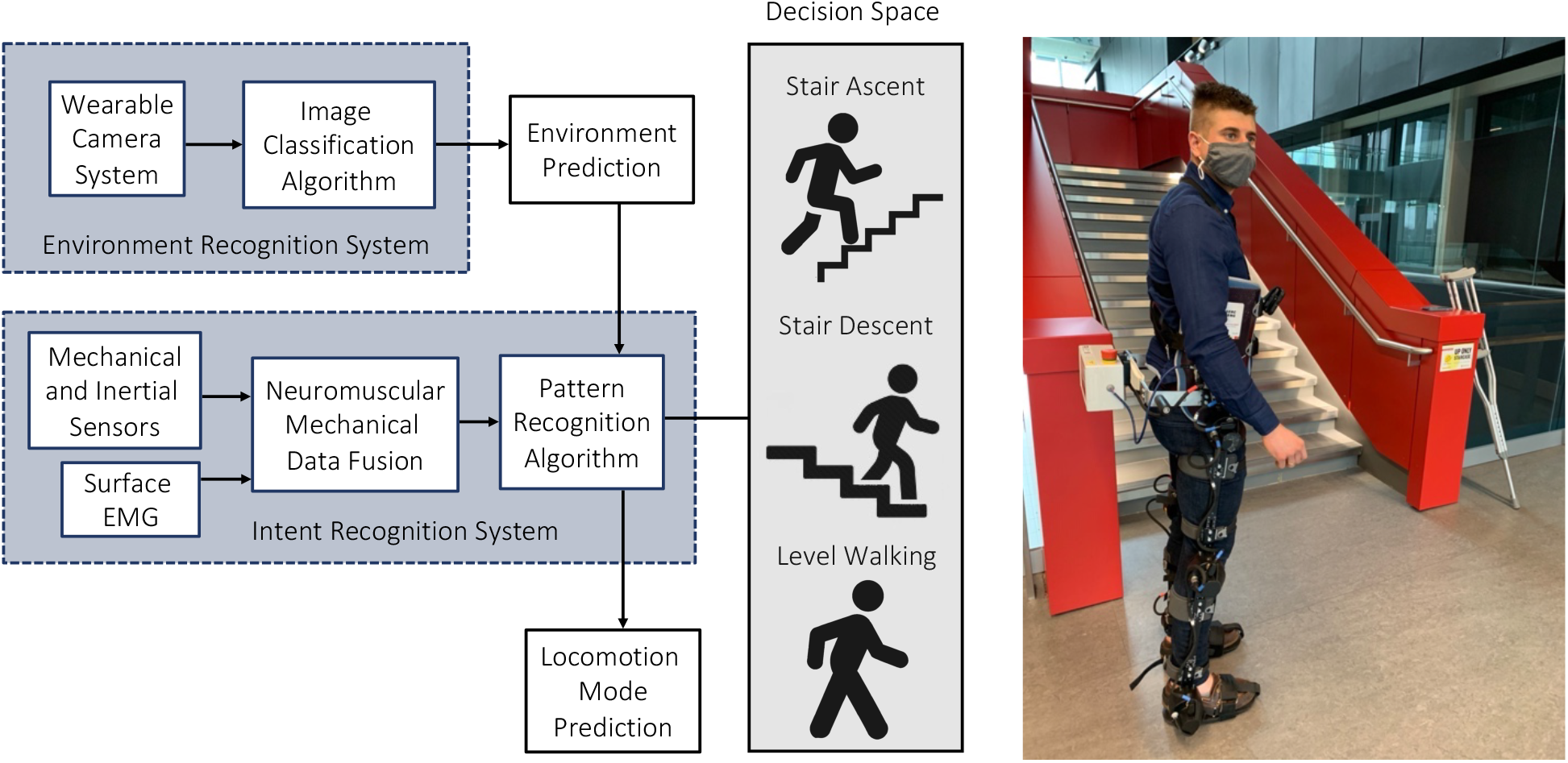
An automated locomotion mode recognition system for robotic prostheses and exoskeletons, also known as an intent recognition system or intelligent high-level controller. These systems can be supplemented with an environment recognition system to forward predict the walking environments prior to physical interaction, therein minimizing the high-level decision space. Note that the photograph (right) is the lead author wearing our robotic exoskeleton.

Mechanical sensors embedded in robotic prostheses and exoskeletons (e.g., potentiometers, pressure sensors, magnetic encoders, and strain gauge load cells) can be used for state estimation by measuring the joint angles and angular velocities, and interaction forces and/or torques between the human and device, and between the device and environment. For example, Varol and colleagues (2010) used onboard mechanical sensors and a Gaussian mixture model classification algorithm for automated intent recognition and high-level control. IMU sensors can measure angular velocities, accelerations, and direction using an onboard gyroscope, accelerometer, and magnetometer, respectively. Although mechanical and inertial sensors allow for fully integrated control systems (i.e., no additional instrumentation or wiring apart from the device need be worn), these sensor technologies can only respond to the user’s movements. In contrast, the electrical potentials of biological muscles, as recorded using surface EMG, precede movement initiation and thus could predict locomotion mode transitions with small prediction horizons. Using only surface EMG and a linear discriminant analysis (LDA) pattern recognition algorithm, Huang et al (2009) differentiated between seven locomotion modes with 90% classification accuracy. In addition to locomotion mode recognition, EMG signals could be used for proportional myoelectric control (Nasr et al., 2021). Although researchers have explored using brain-machine interfaces for direct neural control of robotic prostheses and exoskeletons (i.e., using either implanted or electroencephalography-based systems) (Lebedev and Nicolelis, 2017), these methods are still highly experimental and infrequently used for locomotor applications.

Fusing information from mechanical and/or inertial sensors with surface EMG, known as neuromuscular-mechanical data fusion, has shown to improve the automated locomotion mode recognition accuracies and decision times compared to implementing either system individually (Du et al., 2012; Huang et al., 2011a; 2011b; Krausz and Hargrove, 2021; Liu et al., 2016; Spanias et al., 2015; Wang et al., 2013). Huang et al (2011b) first demonstrated such improvements in classification predictions across six locomotion modes using a support vector machine (SVM) classifier, surface EMG signals, and measured ground reaction forces and torques on a robotic prosthesis. However, neuromuscular-mechanical data are user-dependent, therein requiring time-consuming experiments to amass individual datasets, and surface EMG sensors require calibration and are susceptible to fatigue, motion artifacts, changes in electrode-skin conductivity, and crosstalk between adjacent muscles (Tucker et al., 2015; Young and Ferris, 2017). EMG signals may also be inaccessible in users with high amputations where insufficient neural content is available from the residual limb. Despite the developments in automated intent recognition using mechanical, inertial, and/or neuromuscular signals, further improvements in system performance are desired for safe and robust locomotor control, especially when involving stair environments.

## 2 Literature Review

Taking inspiration from the human vision-locomotor control system, supplementing neuromuscular-mechanical data with information about the oncoming walking environment could improve the automated high-level control performance (see Figure 2). Environment sensing would precede modulation of the user’s muscle activations and/or walking biomechanics (i.e., having an extended prediction horizon), therein allowing for more accurate and real-time transitions between different locomotion modes by minimizing the high-level decision space. During natural human locomotion, the central nervous system acquires state information from biological sensors (e.g., the eyes) through ascending pathways, which are subsequently used to actuate and control the musculoskeletal system through feedforward efferent commands (Patla, 1997; Tucker et al., 2015). However, these control loops are compromised in persons using assistive devices due to limitations in the human-machine data communication. Environment sensing and classification could artificially restore these control loops for automated high-level control. In addition to supplementing the locomotion mode recognition system, environment information could be used to adapt the mid-level reference trajectories (e.g., increasing the actuator joint torques for toe clearance corresponding to an obstacle height) (Zhang et al., 2020), optimal path planning (e.g., identifying opportunities for energy recovery) (Laschowski et al., 2019a; 2021a), and varying foot placement based on the walking surface.

The earliest studies fusing neuromuscular-mechanical data with environment information for prosthetic leg control came from Huang and colleagues (Du et al., 2012; Huang et al., 2011a; Liu et al., 2016; Wang et al., 2013; Zhang et al., 2011). Different walking environments were statistically modelled as prior probabilities using the principle of maximum entropy and incorporated into the discriminant function of an LDA classification algorithm. Rather than assuming equal prior probabilities as usually done, they simulated different walking environments by adjusting the prior probabilities of each class, which allowed their automated locomotion mode recognition system to dynamically adapt to different environments. For instance, when approaching an incline staircase, the prior probability of performing stair ascent would progressively increase while the prior probabilities of other locomotion modes would likewise decrease. Using these adaptive prior probabilities based on the terrain information significantly outperformed (i.e., 95.5% classification accuracy) their locomotion mode recognition system based on neuromuscular-mechanical data alone with equal prior probabilities (i.e., 90.6% accuracy) (Wang et al., 2013). These seminal papers showed 1) how environment information could be incorporated into an automated high-level controller, 2) that including such information could improve the locomotion mode recognition accuracies and decision times, and 3) that the controller could be relatively robust to noisy and imperfect environment predictions such that neuromuscular-mechanical data dominated the high-level decision making (Du et al., 2012; Huang et al., 2011a; Liu et al., 2016; Wang et al., 2013).

Several researchers have explored using wearable radar detectors (Kleiner et al., 2018) and laser rangefinders (Liu et al., 2016; Wang et al., 2013; Zhang et al., 2011) for active environment sensing. Unlike camera-based systems, these sensors circumvent the need for computationally expensive image processing and classification. Radar can uniquely measure distances through non-conducting materials like clothing and are invariant to outdoor lighting conditions and surface textures. Using a leg-mounted radar detector, Herr’s research group (Kleiner et al., 2018) measured stair distances and heights with 1.5 cm and 0.34 cm average accuracies, respectively, up to 6.25 m maximum distances. However, radar reflection signatures usually struggle with source separation of multiple objects and have relatively low resolution. Huang and colleagues (Liu et al., 2016; Wang et al., 2013; Zhang et al., 2011) developed a waist-mounted system with an IMU and laser rangefinder to reconstruct the geometry of the oncoming walking environment between 300-10,000 mm ranges. Environmental features like the terrain height, distance, and slope were used for classification via heuristic rule-based thresholds. The system achieved 98.1% classification accuracy (Zhang et al., 2011). While simple and effective, their system required subject-specific calibration (e.g., the device mounting height) and provided only a single distance measurement.

Compared to radar and laser rangefinders, cameras can provide more detailed information about the field-of-view and detect physical obstacles and terrain changes in peripheral locations (example shown in Figure 3). Most environment recognition systems have used RGB cameras (Da Silva et al., 2020; Diaz et al., 2018; Khademi and Simon, 2019; Krausz and Hargrove, 2015; Laschowski et al., 2019b; 2020b; 2021b; Novo-Torres et al., 2019; Zhong et al., 2020) and/or 3D depth cameras (Hu et al., 2018; Krausz et al., 2015; 2019; Krausz and Hargrove, 2021; Massalin et al., 2018; Varol and Massalin, 2016; Zhang et al., 2019b; 2019c; 2019d, 2020) mounted on the chest (Krausz et al., 2015; Laschowski et al., 2019b; 2020b; 2021b), waist (Khademi and Simon, 2019; Krausz et al., 2019; Krausz and Hargrove, 2021; Zhang et al., 2019d), or lower-limbs (Da Silva et al., 2020; Diaz et al., 2018; Massalin et al., 2018; Varol and Massalin, 2016; Zhang et al., 2019b; 2019c; 2020; Zhong et al., 2020) (see Table 1). Few studies have adopted head-mounted cameras for biomimicry (Novo-Torres et al., 2019; Zhong et al., 2020). Zhong et al (2020) recently compared the effects of different wearable camera positions on classification performance.

**Table 1.**
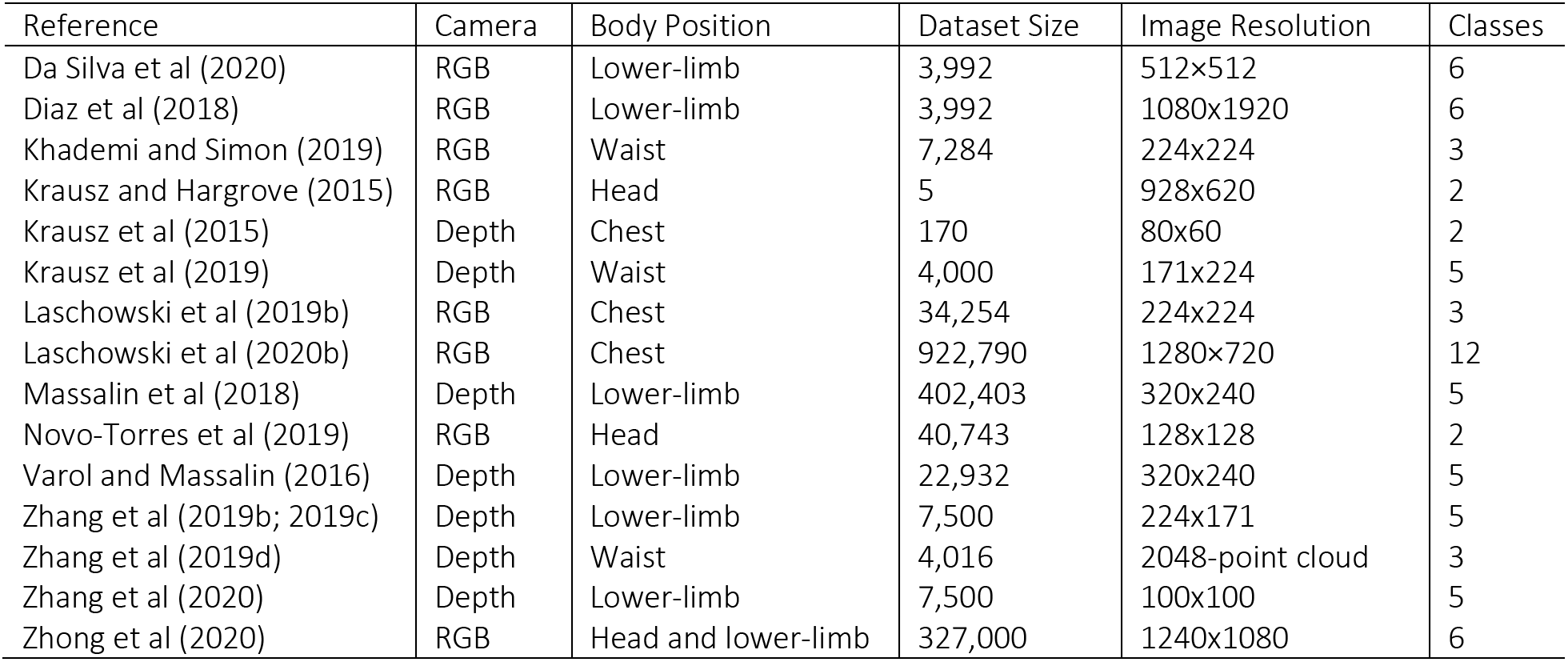
Experimental datasets used for image classification of walking environments for robotic prostheses and exoskeletons. Note that the ExoNet database was published in Laschowski et al (2020b).

| Reference | Camera | Body Position | Dataset Size | Image Resolution | Classes |
| --- | --- | --- | --- | --- | --- |
| Da Silva et al (2020) | RGB | Lower-limb | 3,992 | 512×512 | 6 |
| Diaz et al (2018) | RGB | Lower-limb | 3,992 | 1080x1920 | 6 |
| Khademi and Simon (2019) | RGB | Waist | 7,284 | 224x224 | 3 |
| Krausz and Hargrove (2015) | RGB | Head | 5 | 928x620 | 2 |
| Krausz et al (2015) | Depth | Chest | 170 | 80x60 | 2 |
| Krausz et al (2019) | Depth | Waist | 4,000 | 171x224 | 5 |
| Laschowski et al (2019b) | RGB | Chest | 34,254 | 224x224 | 3 |
| Laschowski et al (2020b) | RGB | Chest | 922,790 | 1280×720 | 12 |
| Massalin et al (2018) | Depth | Lower-limb | 402,403 | 320x240 | 5 |
| Novo-Torres et al (2019) | RGB | Head | 40,743 | 128x128 | 2 |
| Varol and Massalin (2016) | Depth | Lower-limb | 22,932 | 320x240 | 5 |
| Zhang et al (2019b; 2019c) | Depth | Lower-limb | 7,500 | 224x171 | 5 |
| Zhang et al (2019d) | Depth | Waist | 4,016 | 2048-point cloud | 3 |
| Zhang et al (2020) | Depth | Lower-limb | 7,500 | 100x100 | 5 |
| Zhong et al (2020) | RGB | Head and lower-limb | 327,000 | 1240x1080 | 6 |

**Figure 3.**
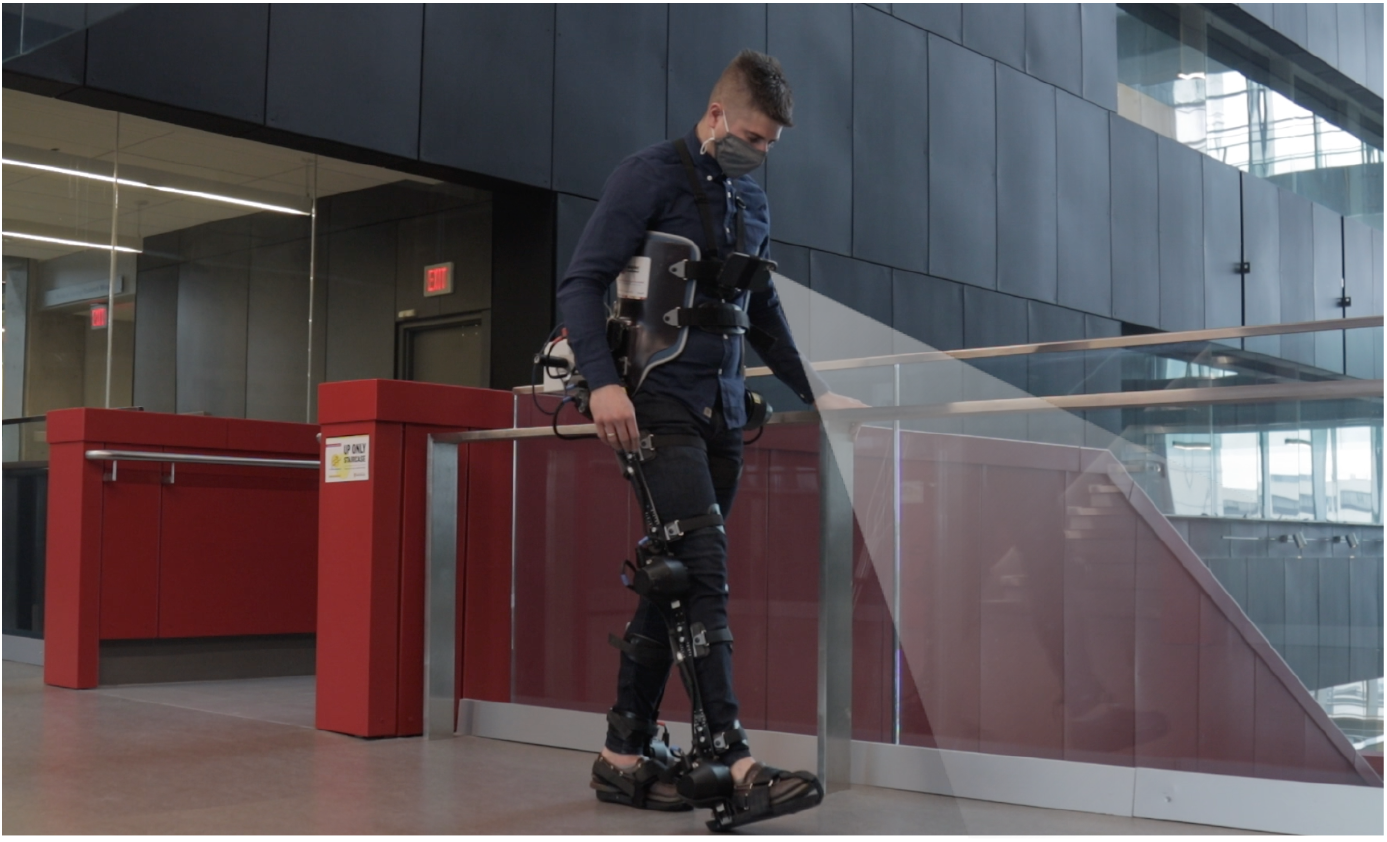
Photograph of the lead author walking with our robotic exoskeleton with camera-based environment sensing superimposed.

Compared to glasses, their leg-mounted camera more accurately detected closer walking environments but struggled with incline stairs, often capturing only 1-2 steps. Although the glasses could detect further-away environments, the head-mounted camera also captured irrelevant features like the sky, which decreased the overall image classification accuracy. The glasses also struggled with detecting decline stairs and had larger standard deviations in classification predictions due to head movement.

For image classification, researchers have traditionally used statistical pattern recognition or machine learning algorithms like support vector machines, which require hand-engineering to develop feature extractors (Da Silva et al., 2020; Diaz et al., 2018; Hu et al., 2018; Krausz et al., 2015; 2019; Krausz and Hargrove, 2021; Massalin et al., 2018; Varol and Massalin, 2016) (see Table 2). Hargrove’s research group (Krausz et al., 2015; 2019) used standard image processing and rule-based thresholds to detect convex and concave edges and vertical and horizontal planes for stair recognition. Although their algorithm achieved 98.8% classification accuracy, the computations were time-consuming (i.e., ~8 seconds/frame) and the system was evaluated using only five sampled images. In another example, Huang and colleagues (Diaz et al., 2018) achieved 86% image classification accuracy across six environment classes using SURF features and a bag-of-words classifier. Varol’s research group used support vector machines for classifying depth images, which mapped extracted features into a high-dimensional space and separated samples into different classes by constructing optimal hyperplanes with maximum margins (Massalin et al., 2018; Varol and Massalin, 2016). Their system achieved 94.1% classification accuracy across five locomotion modes using a cubic kernel SVM and no dimension reduction (Massalin et al., 2018). The average computation time was ~14.9 ms per image. Although SVMs are effective in high-dimensional space and offer good generalization (i.e., robustness to overfitting), these algorithms require manual selection of kernel functions and statistical features, which is time-consuming and suboptimal.

**Table 2.** Previous environment recognition systems that used heuristics, statistical pattern recognition, or support vector machines for image classification of walking environments. Note that each classifier was developed and tested on different image datasets (see Table 1). The computation times are reported per image.

| Reference | Feature Extractor and Classifier | Computing Devices | Test Accuracy (%) | Computation Time (ms) |
| --- | --- | --- | --- | --- |
| Da Silva et al (2020) | Local binary pattern and random forest | NVIDIA Jetson TX2 | 90.0 | 200 |
| Diaz et al (2018) | SURF features and bag-of-words model | Intel Core i7-2600 CPU (3.40GHz) | 86.0 | N/A |
| Krausz and Hargrove (2015) | Hough transform with Gabor filter or canny edge detector | Intel Core i5 | N/A | 8000 |
| Krausz et al (2015) | Heuristic thresholding and edge detector | Intel Core i5 | 98.8 | 200 |
| Krausz et al (2019) | Regions-of-interest and linear discriminate analysis | Intel Core i7-8750H (2.2GHz) | N/A | N/A |
| Massalin et al (2018) | Cubic kernel support vector machine | Intel Core i7-2640M (2.8GHz) | 94.1 | 14.9 |
| Varol and Massalin (2016) | Support vector machine | Intel Core i7-2640M (2.8GHz) | 99.0 | 14.9 |

The latest generation of environment recognition systems has used convolutional neural networks (CNNs) for image classification (Khademi and Simon, 2019; Laschowski et al., 2019b; 2021b; Novo-Torres et al., 2019; Rai and Rombokas, 2018; Zhang et al., 2019b; 2019c; 2019d; 2020; Zhong et al., 2020) (see Table 3 and Figure 4). One of the earliest publications came from Laschowski and colleagues (2019b), who designed and trained a 10-layer convolutional neural network using five-fold cross-validation, which differentiated between three environment classes with 94.9% classification accuracy. The convolutional neural network from Simon’s group (Khademi and Simon, 2019) achieved 99% classification accuracy across three environment classes using transfer learning of pretrained weights. Although CNNs typically outperform SVMs for image classification and circumvent the need for manual feature engineering (LeCun et al., 2015), deep learning requires significant and diverse training data to prevent overfitting and promote generalization. Deep learning has become pervasive in computer vision ever since AlexNet (Krizhevsky et al., 2012) popularized CNNs by winning the 2012 ImageNet challenge with a top-1 accuracy of 63%. ImageNet is an open-source dataset containing ~15 million labelled images and 22,000 classes (Deng et al., 2009). The lack of an open-source, large-scale image dataset of walking environments has impeded the development of environment-adaptive locomotor control systems for robotic prostheses and exoskeletons. To date, each of the aforementioned researchers individually collected experimental data to train their image classification algorithms. These repetitive measurements are time-consuming and inefficient, and individual private datasets have prevented comparisons between classification algorithms from different researchers (Laschowski et al., 2020a).

**Table 3.** Previous environment recognition systems that used convolutional neural networks for image classification of walking environments. Note that each network was trained and tested on different image datasets (see Table 1). The computing hardware were all developed and manufactured by NVIDIA. The number of operations is expressed in multiply-accumulates. The inference times are reported per image.

| Reference | Operations (billions) | Parameters (millions) | Computing Devices | Test Accuracy (%) | Inference Time (ms) |
| --- | --- | --- | --- | --- | --- |
| Khademi and Simon (2019) | 7.7 | 27 | Titan X | 99.6 | 50 |
| Laschowski et al (2019b) | 1.2850 | 4.73 | TITAN Xp | 94.9 | 0.9 |
| Novo-Torres et al (2019) | 0.0011 | 1.13 | Geforce GTX 965M | 90.0 | 5.5 |
| Zhang et al (2019b) | 0.0130 | 0.22 | GeForce GTX 1050 Ti | 96.8 | 3.1 |
| Zhang et al (2019c) | 0.0130 | 0.22 | Quadro P400 | 98.9 | 3.0 |
| Zhang et al (2019d) | 0.0215 | 0.05 | GeForce GTX 1050 Ti | 98.0 | 2.0 |
| Zhang et al (2020) | 0.0130 | 0.22 | Quadro P400 | 96.0 | 3.0 |
| Zhong et al (2020)<sup>†</sup> | 0.0544 | 2.20 | Jetson TX2 | 95.4 | 12.7 |

<sup>†</sup>used MobileNetV2 for feature extraction and a Bayesian neural network (gated recurrent unit) for the environment classification.

**Figure 4.**
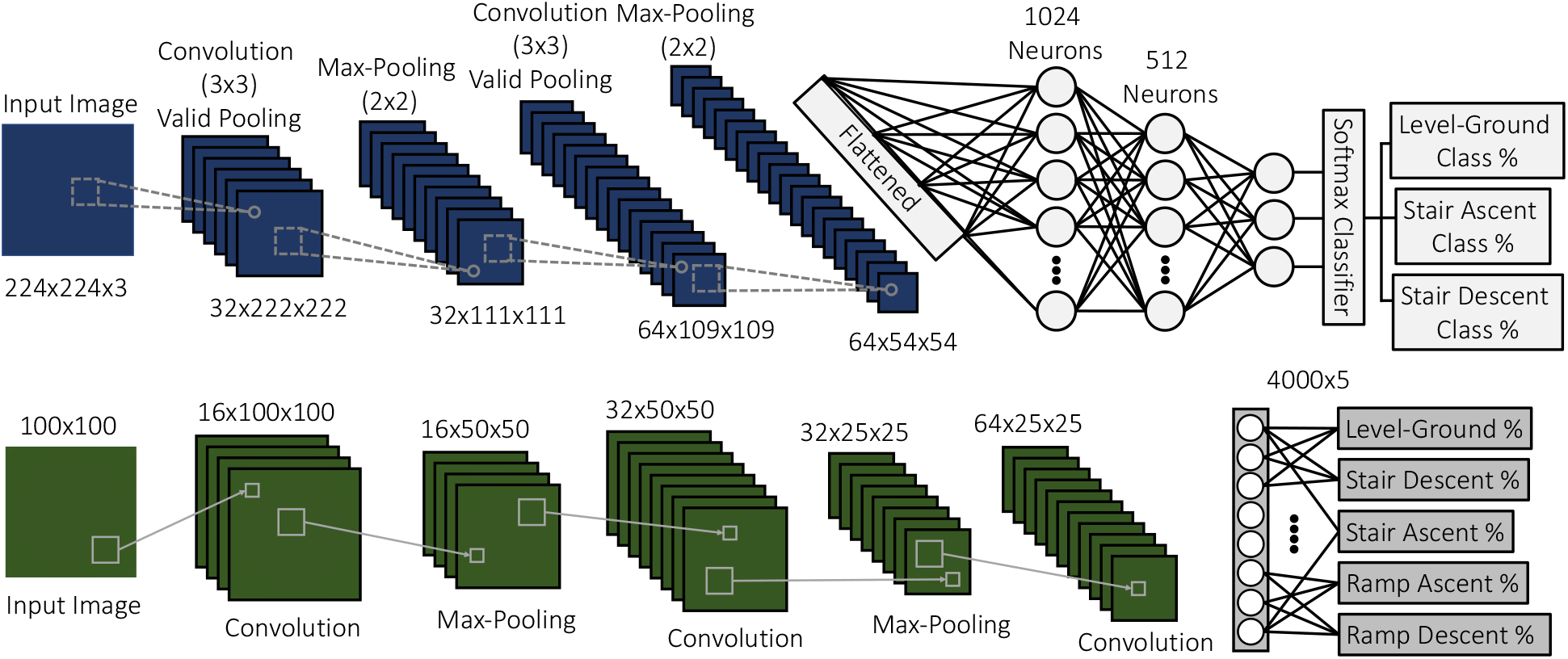
Examples of convolutional neural networks previously used for image classification of walking environments. The top and bottom schematics were adapted from Dan Simon (Cleveland State University, USA) and Chenglong Fu (Southern University of Science and Technology, China), respectively.

Motivated by these limitations in the literature, we developed and evaluated an advanced environment classification system that uses computer vision and deep learning. In this study, we 1) reviewed the development of the “ExoNet” database – the largest and most diverse open-source dataset of wearable camera images of human walking environments; 2) trained and tested over a dozen state-of-the-art deep convolutional neural networks on the ExoNet database for large-scale image classification of the different walking environments; and 3) quantitatively compared the benchmarked CNN architectures and their environment classification predictions using an operational metric called “NetScore”, which balances the image classification accuracy with the computational and memory storage requirements (i.e., important for onboard real-time inference). The overall objective of this research is to support the development of next-generation environment-adaptive locomotion mode recognition systems for robotic prostheses and exoskeletons.

## 3 Materials and Methods

### 3.1 Environment Sensing

One participant without wearing an assistive device was instrumented with a lightweight smartphone camera system (iPhone XS Max) (see Figure 5). Compared to lower-limb systems (Da Silva et al., 2020; Diaz et al., 2018; Hu et al., 2018; Kleiner et al., 2018; Massalin et al., 2018; Rai and Rombokas, 2018; Varol and Massalin, 2016; Zhang et al., 2011; 2019b; 2019c; 2020), our chest-mounted camera can provide more stable video recording and allow users to wear pants and dresses without obstructing the sampled field-of-view. The chest-mount height was ~1.3 m from the ground when the participant stood upright. The smartphone has two 12-megapixel RGB rear-facing cameras and one 7-megapixel front-facing camera. The front and rear cameras provide 1920×1080 and 1280×720 video recording at 30 Hz, respectively. The smartphone weighs 0.21 kg and has an onboard rechargeable lithium-ion battery, 512-GB of memory storage, and a 64-bit ARM-based integrated circuit (Apple A12 Bionic) with a six-core CPU and four-core GPU; these hardware specifications can theoretically support onboard deep learning for real-time environment classification. The relatively lightweight and unobtrusive nature of the wearable camera system allowed for unimpeded human locomotion. Ethical review and approval were not required for this research in accordance with the University of Waterloo Office of Research Ethics.

Whereas most environment recognition systems have been limited to controlled indoor environments and/or prearranged walking circuits (Du et al., 2012; Hu et al., 2018; Khademi and Simon, 2019; Kleiner et al., 2018; Krausz et al., 2015; 2019; Krausz and Hargrove, 2021; Liu et al., 2016; Rai and Rombokas, 2018; Wang et al., 2013; Zhang et al., 2011; 2019b; 2019c; 2019d), our participant walked around unknown outdoor and indoor real-world environments while collecting images with occlusions and intraclass variations (see Figure 6). We collected data at various times throughout the day to include different lighting conditions. Similar to human gaze fixation during walking (Li et al., 2019), the field-of-view was 1-5 m ahead of the participant, therein showing the oncoming walking environment rather than the ground directly underneath the subject’s feet. This operating range allowed for detecting physical obstacles and terrain changes within several walking strides; research has shown that lower-limb amputees tend to allocate higher visual attention to oncoming terrain changes compared to healthy controls (Li et al., 2019). Our camera’s pitch angle slightly differed between data collection sessions. Images were sampled at 30 Hz with 1280×720 resolution. We recorded over 52 hours of video, amounting to ~5.6 million images. The same environment was never sampled twice to maximize diversity in the dataset. Images were collected during the summer, fall, and winter seasons to capture different weathered surfaces like snow, grass, and multicolored leaves. The image database, which we named “ExoNet”, was uploaded to the IEEE DataPort repository and is publicly available for download (Laschowski et al., 2020b). The file size of the uncompressed videos is ~140 GB.

**Figure 5.**
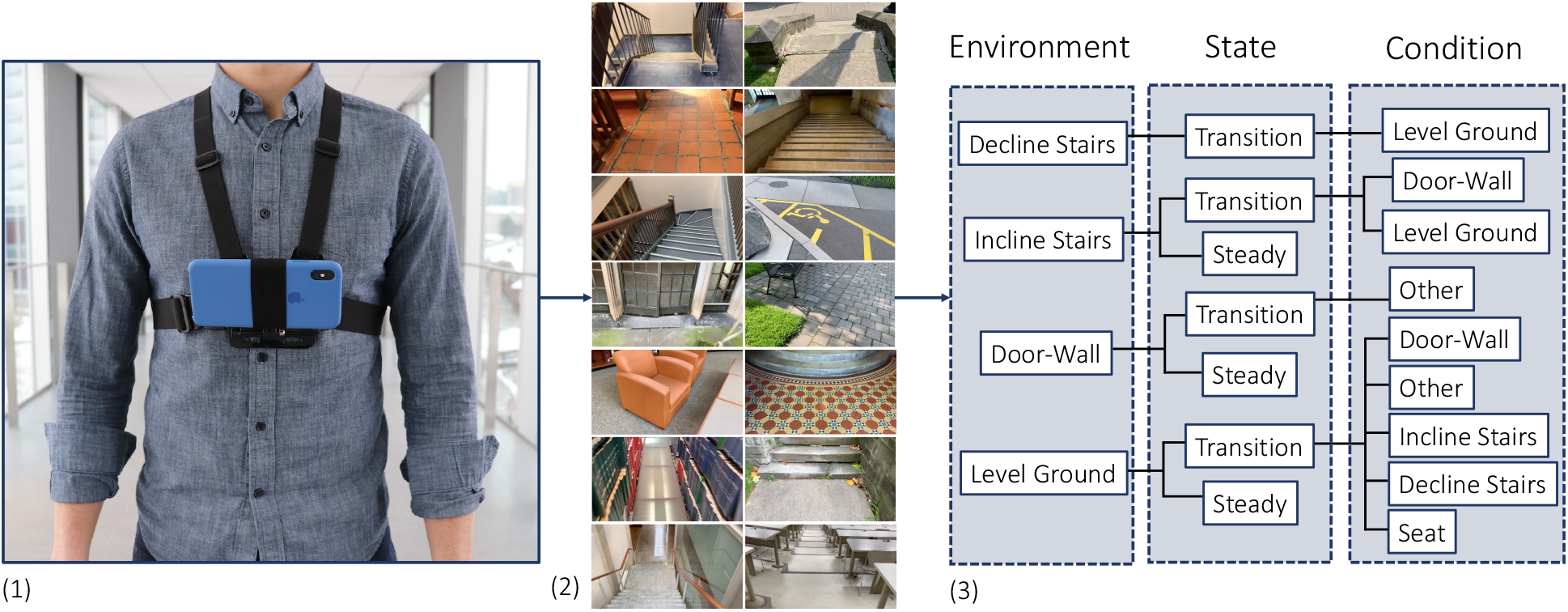
Development of the “ExoNet” database, including (1) photograph of the wearable camera system used for large-scale data collection; (2) examples of high-resolution RGB images of walking environments; and (3) schematic of the 12-class hierarchical labelling architecture.

**Figure 6.**
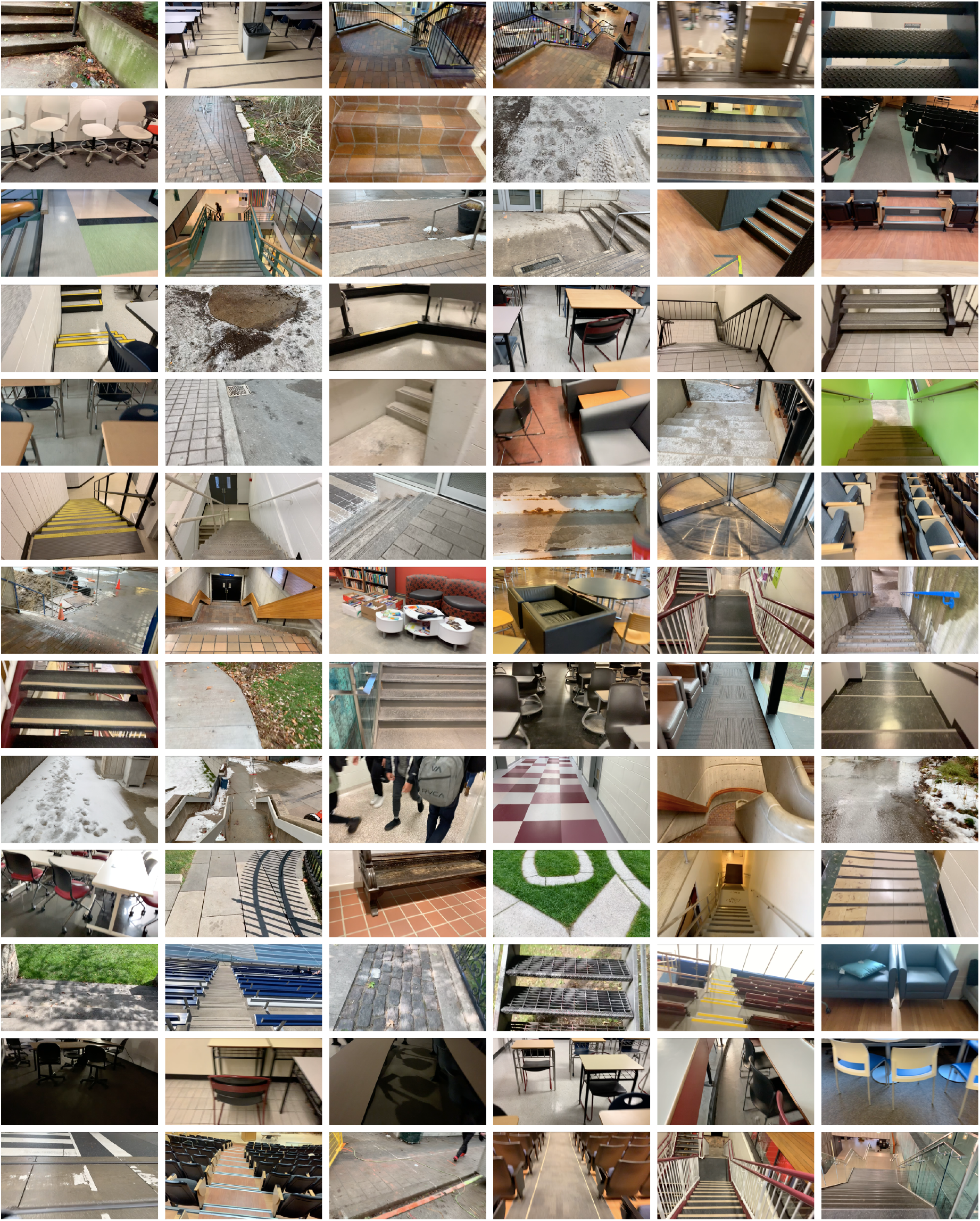
Examples of the wearable camera images of indoor and outdoor real-world walking environments in the ExoNet database. Images were collected at various times throughout the day and across different seasons (i.e., summer, fall, and winter).

Given the subject’s self-selected walking speed, there were relatively minimal differences between consecutive images sampled at 30 Hz. We therefore downsampled and labelled the images at 5 frames/second to minimize the demands of manual annotation and increase the diversity in image appearances. However, for online environment-adaptive control of robotic leg prostheses and exoskeletons, higher sampling rates would be more advantageous for accurate and robust automated locomotion mode transitions. Similar to the ImageNet dataset (Deng et al., 2009), the ExoNet database was human-annotated using a hierarchical labelling architecture. Images were mainly labelled according to common high-level locomotion modes, rather than a purely computer vision perspective. For instance, images of level-ground environments showing either pavement or grass were not differentiated since both surfaces would be assigned the same level-ground walking locomotion mode. In contrast, computer vision researchers might label these different surface textures as separate classes.

Approximately 923,000 images were manually annotated using a 12-class hierarchical labelling architecture (see Figure 5). The dataset included: 31,628 images of “incline stairs transition wall/door” (I-T-W); 11,040 images of “incline stairs transition level-ground” (I-T-L); 17,358 images of “incline stairs steady” (I-S); 28,677 images of “decline stairs transition level-ground” (D-T-L); 19,150 images of “wall/door transition other” (W-T-O); 36,710 images of “wall/door steady” (W-S); 379,199 images of “level-ground transition wall/door” (L-T-W); 153,263 images of “level-ground transition other” (L-T-O); 26,067 images of “level-ground transition incline stairs” (L-T-I); 22,607 images of “level-ground transition decline stairs” (L-T-D); 119,515 images of “level-ground transition seats” (L-T-E); and 77,576 images of “level-ground steady” (L-S). These class labels were chosen and assigned post hoc to capture the different walking environments from the data collection. In comparison, Novo-Torres et al (2019) simultaneously sampled and labelled their images using a portable keyboard during data collection. Similar to previous research (Liu et al., 2016; Wang et al., 2013; Zhang et al., 2011), we included an “other” class to maintain the image classification performance when unlabeled environments and/or objects like pedestrians, cars, and bicycles were observable.

Taking inspiration from (Du et al., 2012; Huang et al., 2011a; Khademi and Simon, 2019; Liu et al., 2016; Wang et al., 2013), our labelling architecture included both “steady” (S) and “transition” (T) states (see Figure 7). A steady state describes an environment where an exoskeleton or prosthesis user would continue to perform the same locomotion mode (e.g., an image showing only level-ground terrain). In contrast, a transition state describes an environment where an exoskeleton or prosthesis high-level controller might switch between locomotion modes (e.g., an image showing both level-ground terrain and incline stairs). Manually labelling these transition states was relatively subjective. For instance, an image showing level-ground terrain was labelled “level-ground transition incline stairs” (L-T-I) when an incline staircase was approximately within the sampled field-of-view. Similar labelling principles were applied to transitions to other conditions. Although the ExoNet database was labelled by one designated researcher, consistently determining the exact video frame where an environment would switch between steady and transition states was challenging; Huang et al (2011a) reported experiencing similar difficulties.

**Figure 7.**
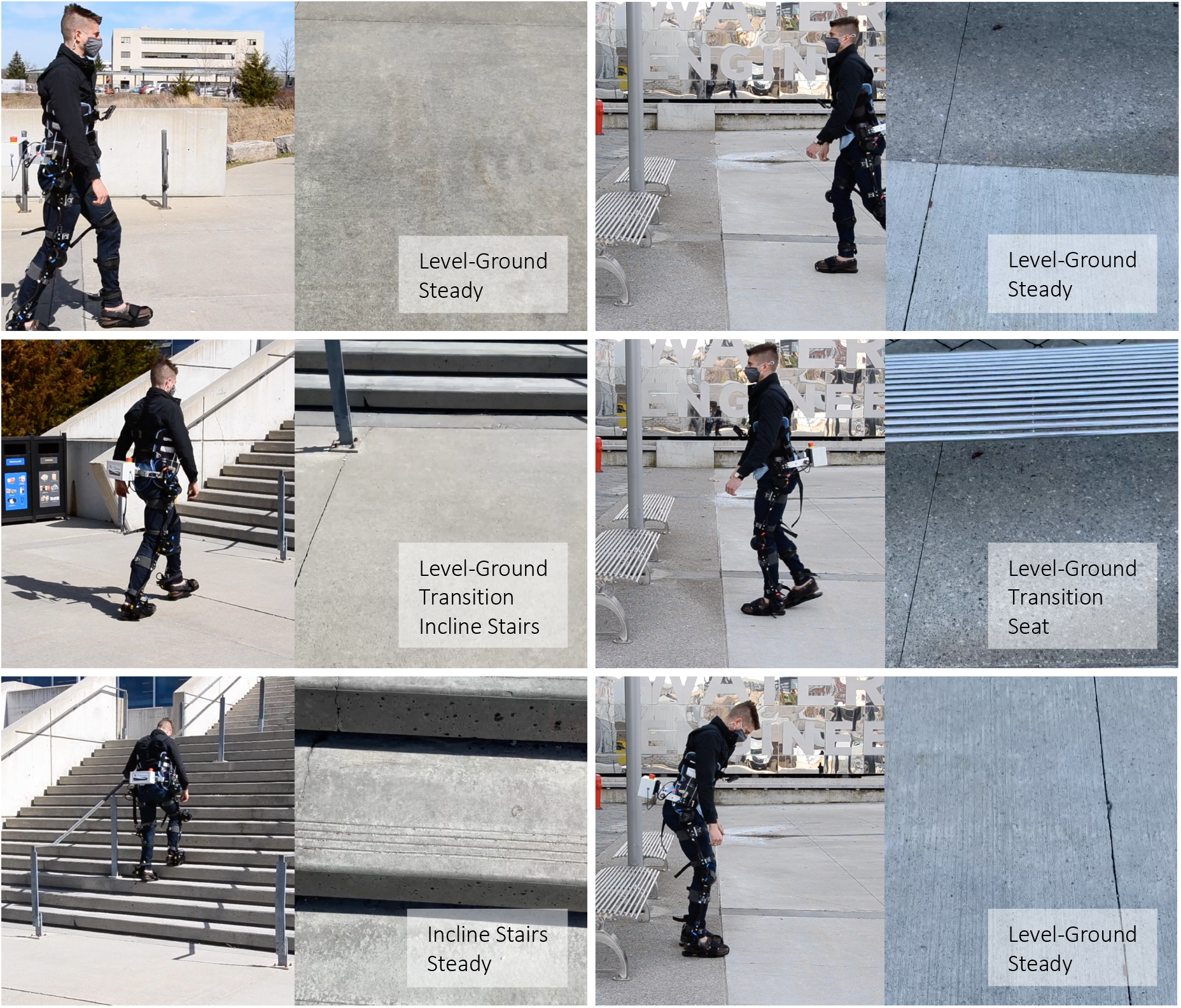
Examples of both “steady” and “transition” states in the ExoNet hierarchical labelling architecture. The top and bottom rows are labelled as steady states and the middle row is labelled a transition state. For each column, the left images show the lead author walking with our robotic exoskeleton and the right images show the concurrent field-of-view of the wearable camera system (i.e., what the exoskeleton sees).

### 3.2 Environment Classification

We used convolutional neutral networks for the image classification. Unlike previous studies that used support vector machines that required hand-engineering (Massalin et al., 2018; Varol and Massalin, 2016), deep learning replaces manually extracted features with multilayer networks that can automatically and efficiently learn the optimal image features from the training data. Generally speaking, a CNN architecture contains multiple stacked convolutional and pooling layers with decreasing spatial resolutions and increasing number of feature maps. Starting with an input 3D tensor, the convolutional layers perform convolution operations (i.e., dot products) between the inputs and convolutional filters. The first few layers extract relatively general features, like edges, while deeper layers learn more problem-dependent features. The resulting feature maps are passed through a nonlinear activation function, typically a rectified linear unit (ReLU). The pooling layers spatially downsample the feature maps to reduce the computational effort by aggregating neighboring elements using either maximum or average values. The CNN architecture concludes with one or more fully connected layers and a softmax loss function, which estimates the probability distribution (i.e., scores) of each labelled class. Previous CNN-based environment recognition systems (see Table 3) have used supervised learning such that the differences between the predicted and labelled class scores are computed and the learnable network parameters (i.e., weights) are optimized to minimize the loss function via backpropagation and stochastic gradient descent. After training, the CNN performs inference on previously unseen data to evaluate the generalizability of the learned parameters. These artificial neural networks are loosely inspired by biological neural networks.

We used TensorFlow version 2.3 and the Keras high-level API to train and test over a dozen state-of-the-art deep convolutional neutral networks on the ExoNet database for large-scale image classification, including: EfficientNetB0 (Tan and Le, 2019); InceptionV3 (Szegedy et al., 2015); MobileNet (Howard et al., 2017); MobileNetV2 (Sandler et al., 2018); VGG16 and VGG19 (Simonyan and Zisserman, 2014); Xception (Chollet, 2016); ResNet50, ResNet101, and ResNet152 (He et al., 2015); and DenseNet121, DenseNet169, and DenseNet201 (Huang et al., 2017). During data preprocessing, the images were cropped to an aspect ratio of 1:1 and downsampled to 256×256 using bilinear interpolation. Random crops of 224×224 were used as inputs to the neural networks; this method of data augmentation helped further increase the sample diversity. The final densely connected layer of each CNN architecture was modified by setting the number of output channels equal to the number of environment classes in the ExoNet database. We used a softmax loss function to predict the individual class scores. The labelled images were split into training (89.5%), validation (3.5%), and testing (7%) sets, the proportions of which are consistent with ImageNet (Deng et al., 2009), which is of comparable size. We experimented with transfer learning of pretrained weights from ImageNet but found no additional performance benefit.

Dropout regularization was applied before the final dense layer to prevent overfitting during training such that the learnable weights were randomly dropped (i.e., activations set to zero) during each forward pass at a rate of 0.5. Images were also randomly flipped horizontally during training to increase stochasticity and promote generalization. We trained each CNN architecture for 40 epochs using a batch size and initial learning rate of 128 and 0.001, respectively; these hyperparameters were experimentally tuned to maximize the performance on the validation set (see Figure 8). We explored different combinations of batch sizes of 32, 64, 128, and 256; epochs of 20, 40, and 60; dropout rates of 0, 0.2, 0.5; and initial learning rates of 0.01, 0.001, 0.0001, and 0.00001. The learning rate was reduced during training using a cosine weight decay schedule (Loshchilov and Hutter, 2016). We calculated the sparse categorical cross-entropy loss between the labelled and predicted classes and used the Adam optimization algorithm (Kingma and Ba, 2015), which computes backpropagated gradients using momentum and an adaptive learning rate, to update the learnable weights and minimize the loss function. During testing, we used a single central crop of 224×224. Training and inference were both performed on a Tensor Processing Unit (TPU) version 3-8 by Google Cloud; these customized chips allow for accelerated CNN computations (i.e., matrix multiplications and additions) compared to traditional computing devices.

**Figure 8.**
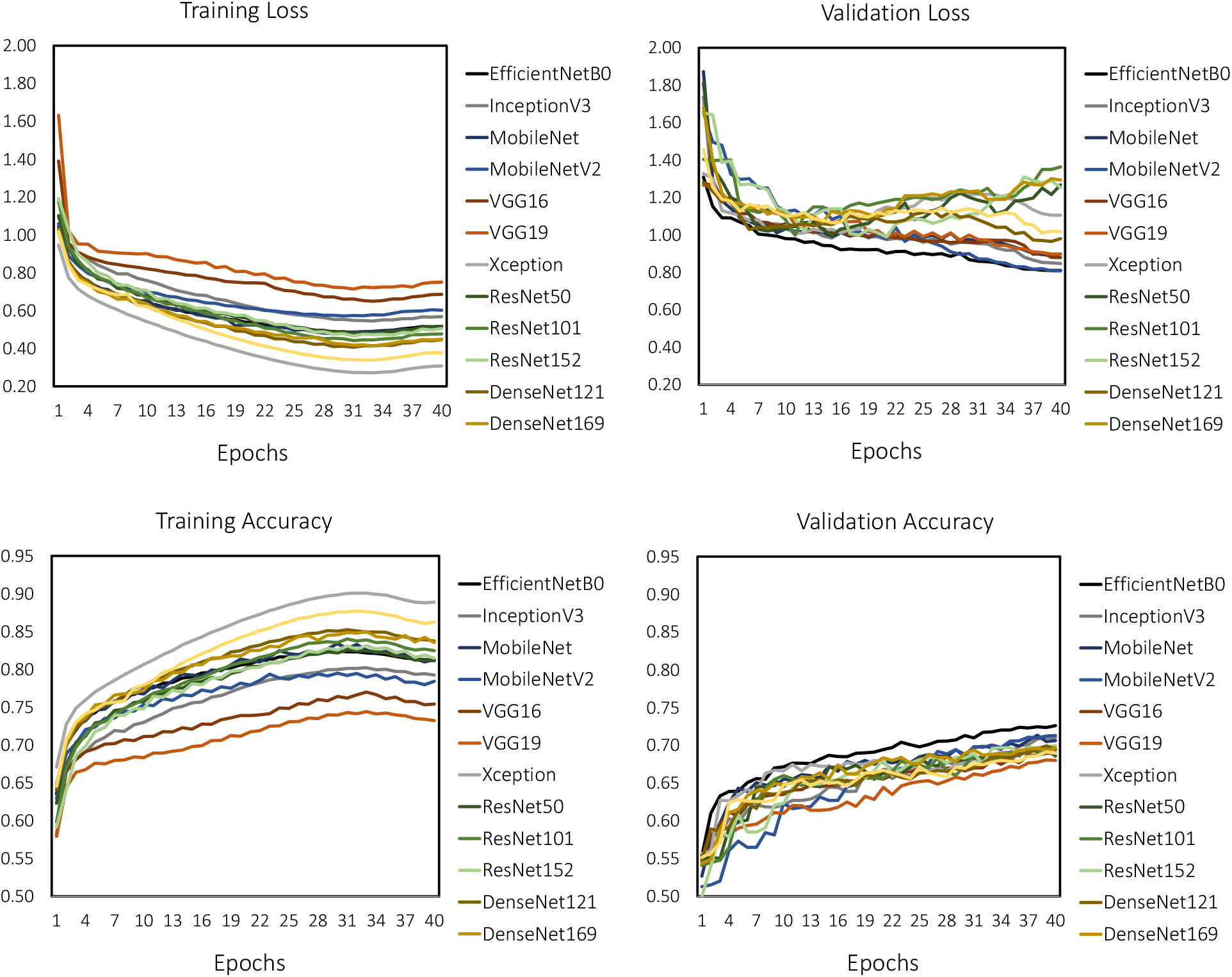
The loss and image classification accuracies during training and validation on the ExoNet database using state-of-the-art deep convolutional neural networks, including: EfficientNetB0, InceptionV3, MobileNet, MobileNetV2, VGG16, VGG19, Xception, ResNet50, ResNet101, ResNet152, DenseNet121, DenseNet169, and DenseNet201.

### 3.3 NetScore Evaluation

To facilitate onboard real-time inference for environment-adaptative control of robotic leg prostheses and exoskeletons, the ideal convolutional neural network would achieve high classification accuracy with minimal parameters, computing operations, and inference time. Motivated by these design principles, we quantitatively evaluated and compared the benchmarked CNN architectures (Ɲ) and their environment classification predictions on the ExoNet database using an operational metric called “NetScore” (Wong, 2018):

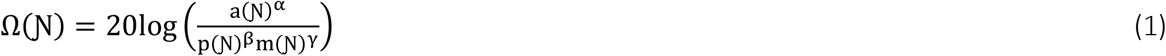

where a(Ɲ) is the image classification accuracy during inference (0-100%), p(Ɲ) is the number of parameters expressed in millions, m(Ɲ) is the number of multiply–accumulates expressed in billions, and α, β, and γ are coefficients that control the effects of the classification accuracy, and the architectural and computational complexities on the NetScore (Ω), respectively. We set the coefficients to {α = 2, β = 0.5, γ = 0.5} to better emphasize the classification accuracy while partially considering the parameters and computing operations, since neural networks with low accuracy are less practical, regardless of the size and speed. Note that the NetScore does not explicitly account for inference time. The number of parameters p(Ɲ) and multiply–accumulates m(Ɲ) are assumed to be representative of the architectural and computational complexities, respectively, both of which are inversely proportional to the NetScore.

## 4 Results

Table 4 summarizes the benchmarked CNN architectures (i.e., the number of parameters and computing operations) and their environment classification performances on the ExoNet database (i.e., prediction accuracies and inference times), including the NetScores. The EfficientNetB0 network achieved the highest image classification accuracy (Ca) during inference (i.e., 73.2% accuracy), that being the percentage of true positives (i.e., 47,265 images) out of the total number of images in the testing set (i.e., 64,568 images) 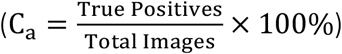. In contrast, the VGG19 network produced the least accurate predictions, with an overall image classification accuracy of 69.2%. The range of accuracies across the different CNN architectures was thus relatively small (i.e., maximum arithmetic difference of 4 percentage points). We observed relatively weak correlations between both the number of parameters (i.e., Pearson r = −0.3) and computing operations (i.e., Pearson r = −0.59) and the classification accuracies on the ExoNet database across the different CNN architectures.

**Table 4.**
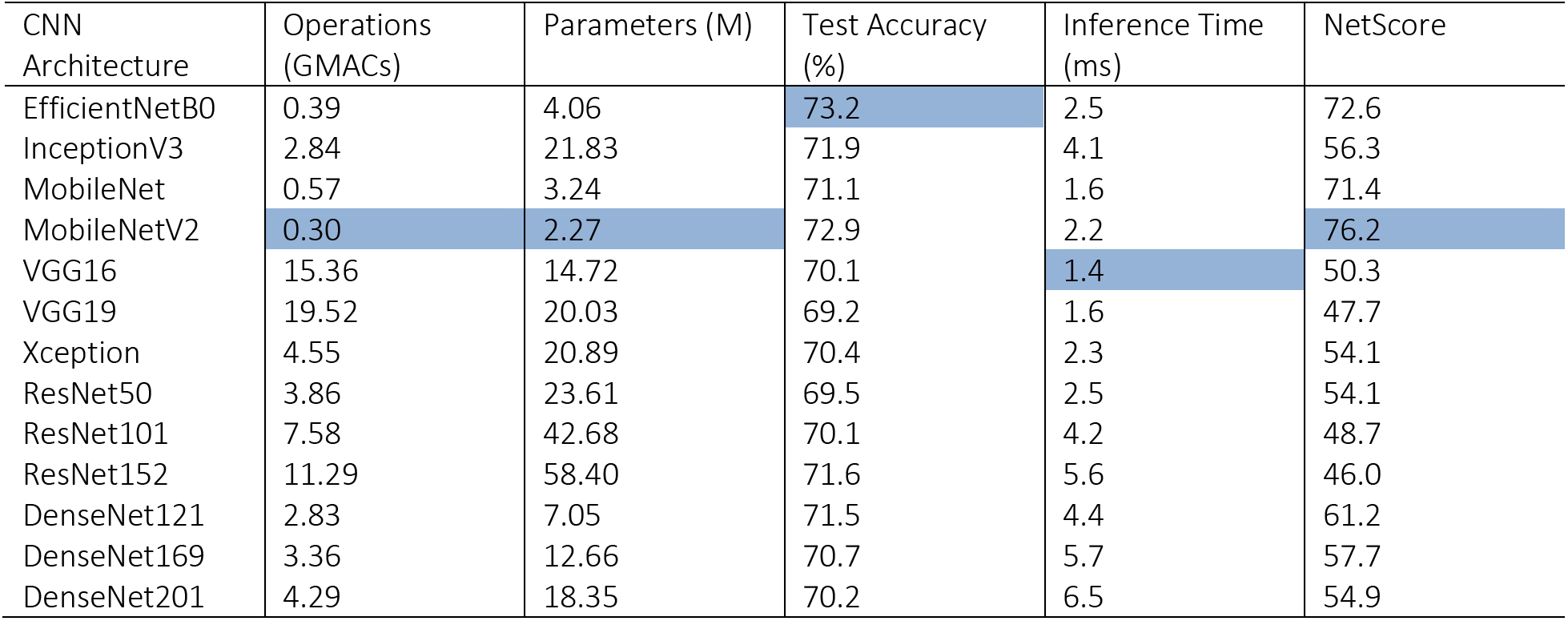
The benchmarked CNN architectures and their environment classification performances during inference on the ExoNet database. The test accuracies, parameters, and computing operations are expressed in percentages (0-100%), millions of parameters (M), and billions of multiply-accumulates (GMACs), respectively. Training and inference were both performed on a Google Cloud TPU. The EfficientNetB0 network achieved the highest test accuracy, VGG16 the fastest inference time, and MobileNetV2 the best NetScore and least number of parameters and computing operations (highlighted below).

Although the VGG16 and VGG19 networks involved the largest number of computations – i.e., ~15.4 and ~19.5 billion multiply-accumulates, respectively – they resulted in the fastest inference times (i.e., on average 1.4 ms and 1.6 ms per image). For comparison, the DenseNet201 network had 72.1% and 78% fewer operations than VGG16 and VGG19, respectively, but was 364% and 306% slower. These results concur with Ding et al (2021), who recently showed 1) that the number of computing operations does not explicitly reflect the actual inference speed, and 2) that VGG-style architectures can run faster and more efficiently on CNN computing devices compared to more complicated architectures like DenseNets due to their relatively simple designs (i.e., consisting of basic convolutions and ReLU activations). Our inference times were calculated on the Cloud TPU using a batch size of 8. Note that the relative inference speeds between the benchmarked CNN architectures (i.e., their ordering from fastest to slowest) may differ across different computing devices given that some platforms are designed to accelerate certain operations better than others (e.g., cloud computing vs. those designed for mobile and embedded systems).

Tables 5–17 show the multiclass confusion matrix for each CNN architecture. The matrix columns and rows are the predicted and labelled classes, respectively. The diagonal elements are the classification accuracies for each environment class during inference, known as true positives, and the nondiagonal elements are the misclassification percentages; darker shades represent higher classification accuracies. Despite slight numerical differences, most of the benchmarked CNN architectures displayed a similar interclass trend. The networks most accurately predicted the “level-ground transition wall/door” (L-T-W) class (i.e., average Ca = 84.3% ± 2.1%), followed by the “level-ground steady” (L-S) class with an average accuracy of 77.3% ± 2.1% and the “decline stairs transition level-ground” (D-T-L) class with an average accuracy of 76.6% ± 2.4%. These results could be attributed to the class imbalances among the training data (i.e., there were significantly more images of L-T-W environments compared to other classes). However, some classes with limited images showed relatively good classification performance. For instance, the “incline stairs transition level-ground” (I-T-L) class comprised only 1.2% of the ExoNet database but achieved 71.2% ± 5.1% average classification accuracy. The least accurate predictions were for the environment classes that contained “other” features – i.e., the “wall/door transition other” (W-T-O) class with an average accuracy of 38.3% ± 3.5% and the “level-ground transition other” (L-T-O) class with an average accuracy of 46.8% ± 2.4%; these results could be attributed to the increased noise and randomness of the environments and/or objects in the images.

**Table 5.**
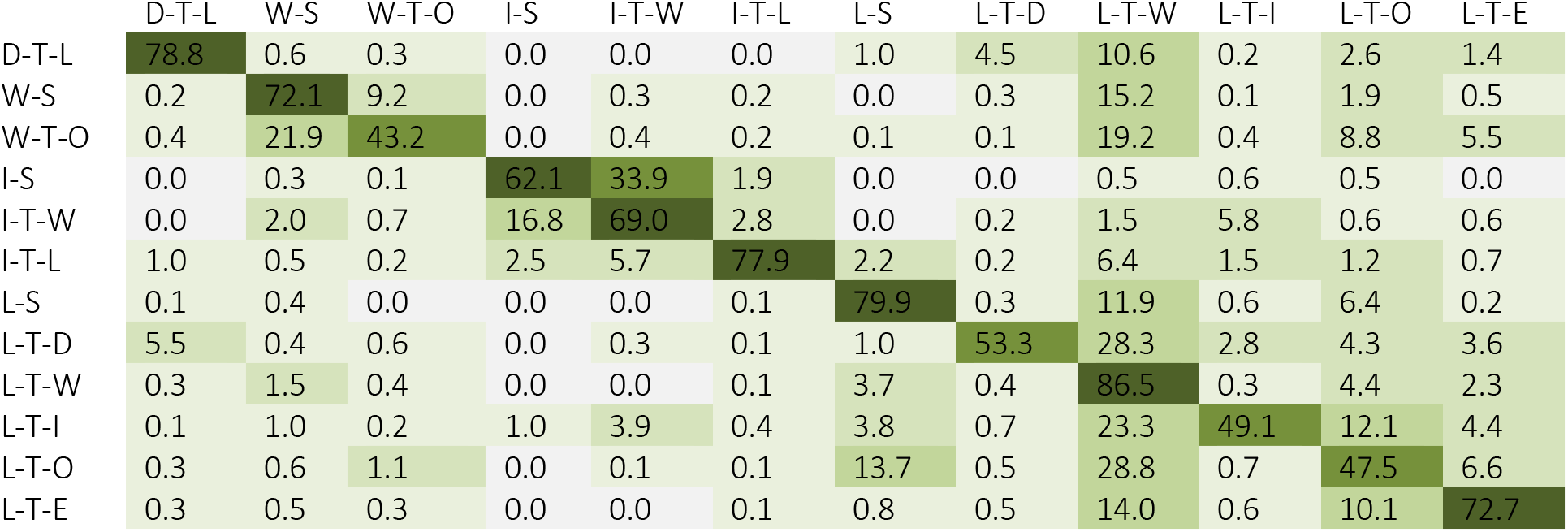
The multiclass confusion matrix for EfficientNetB0 showing the image classification accuracies (%) during inference on the ExoNet database. The columns and rows are the predicted and labelled classes, respectively.

**Table 6.** The multiclass confusion matrix for InceptionV3 showing the image classification accuracies (%) during inference on the ExoNet database. The columns and rows are the predicted and labelled classes, respectively.

|  | D-T-L | W-S | W-T-O | I-S | I-T-W | I-T-L | L-S | L-T-D | L-T-W | L-T-I | L-T-O | L-T-E |
| --- | --- | --- | --- | --- | --- | --- | --- | --- | --- | --- | --- | --- |
| D-T-L | 77.3 | 0.8 | 0.3 | 0.0 | 0.1 | 0.4 | 1.1 | 4.3 | 11.0 | 0.2 | 2.6 | 1.9 |
| W-S | 0.3 | 70.6 | 10.9 | 0.0 | 0.2 | 0.1 | 0.0 | 0.2 | 16.3 | 0.1 | 1.0 | 0.3 |
| W-T-O | 0.7 | 22.7 | 38.4 | 0.0 | 0.3 | 0.0 | 0.2 | 0.5 | 21.0 | 0.5 | 12.0 | 3.7 |
| I-S | 0.0 | 0.8 | 0.1 | 66.6 | 25.9 | 3.3 | 0.8 | 0.0 | 1.0 | 1.0 | 0.3 | 0.3 |
| I-T-W | 0.1 | 2.0 | 0.9 | 15.1 | 68.7 | 3.2 | 0.0 | 0.0 | 2.9 | 5.3 | 1.0 | 0.8 |
| I-T-L | 0.3 | 1.3 | 0.2 | 2.5 | 3.9 | 72.4 | 4.0 | 0.2 | 10.3 | 2.4 | 1.0 | 1.5 |
| L-S | 0.2 | 0.1 | 0.0 | 0.0 | 0.0 | 0.1 | 78.6 | 0.2 | 12.5 | 0.3 | 7.4 | 0.5 |
| L-T-D | 6.1 | 0.6 | 0.4 | 0.0 | 0.4 | 0.1 | 1.1 | 53.3 | 28.2 | 1.1 | 5.1 | 3.6 |
| L-T-W | 0.3 | 1.9 | 0.5 | 0.0 | 0.1 | 0.1 | 4.2 | 0.5 | 85.5 | 0.4 | 4.3 | 2.2 |
| L-T-I | 0.0 | 0.9 | 0.4 | 0.5 | 4.0 | 0.5 | 4.4 | 1.4 | 24.9 | 47.6 | 10.8 | 4.5 |
| L-T-O | 0.4 | 0.5 | 1.1 | 0.0 | 0.1 | 0.0 | 14.5 | 0.5 | 30.7 | 0.6 | 46.2 | 5.3 |
| L-T-E | 0.3 | 0.5 | 0.6 | 0.0 | 0.1 | 0.1 | 0.9 | 0.4 | 16.1 | 0.7 | 9.7 | 70.6 |

**Table 7.**
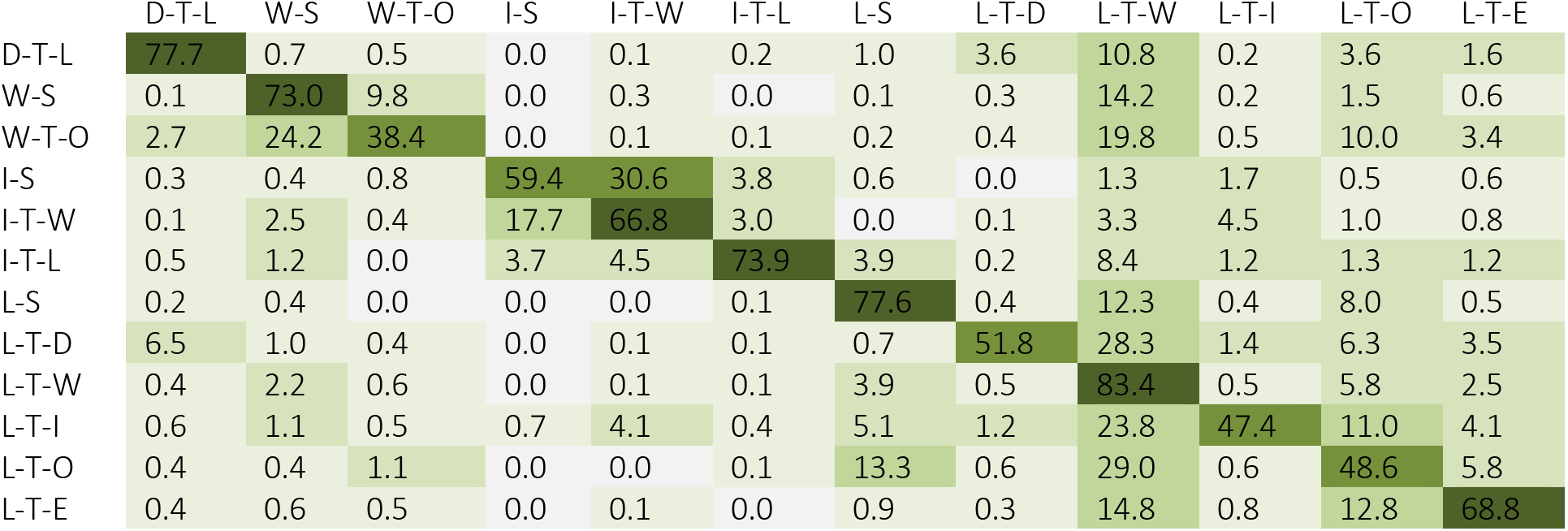
The multiclass confusion matrix for MobileNet showing the image classification accuracies (%) during inference on the ExoNet database. The columns and rows are the predicted and labelled classes, respectively.

**Table 8.** The multiclass confusion matrix for MobileNetV2 showing the image classification accuracies (%) during inference on the ExoNet database. The columns and rows are the predicted and labelled classes, respectively.

|  | D-T-L | W-S | W-T-O | I-S | I-T-W | I-T-L | L-S | L-T-D | L-T-W | L-T-I | L-T-O | L-T-E |
| --- | --- | --- | --- | --- | --- | --- | --- | --- | --- | --- | --- | --- |
| D-T-L | 78.3 | 0.6 | 0.4 | 0.0 | 0.1 | 0.0 | 0.9 | 5.0 | 9.9 | 0.2 | 2.9 | 1.8 |
| W-S | 0.2 | 73.4 | 10.1 | 0.0 | 0.3 | 0.3 | 0.1 | 0.4 | 13.4 | 0.2 | 1.2 | 0.5 |
| W-T-O | 0.6 | 24.2 | 41.5 | 0.0 | 0.4 | 0.0 | 0.0 | 0.6 | 18.0 | 0.6 | 9.5 | 4.6 |
| I-S | 0.0 | 0.4 | 0.1 | 64.7 | 28.9 | 2.6 | 0.4 | 0.0 | 0.9 | 1.3 | 0.1 | 0.6 |
| I-T-W | 0.0 | 2.0 | 0.9 | 18.5 | 66.2 | 3.3 | 0.0 | 0.1 | 2.7 | 4.9 | 0.5 | 1.0 |
| I-T-L | 0.2 | 0.7 | 0.0 | 2.2 | 7.9 | 73.6 | 3.4 | 0.2 | 7.9 | 1.7 | 1.0 | 1.3 |
| L-S | 0.1 | 0.3 | 0.0 | 0.0 | 0.0 | 0.1 | 79.2 | 0.1 | 11.9 | 0.2 | 7.7 | 0.4 |
| L-T-D | 5.5 | 0.7 | 0.4 | 0.0 | 0.0 | 0.1 | 1.0 | 53.8 | 28.2 | 1.1 | 5.5 | 3.7 |
| L-T-W | 0.3 | 1.9 | 0.5 | 0.0 | 0.0 | 0.1 | 3.9 | 0.4 | 86.5 | 0.3 | 4.1 | 2.0 |
| L-T-I | 0.2 | 1.2 | 0.1 | 0.1 | 3.9 | 0.4 | 4.1 | 1.0 | 26.1 | 48.4 | 9.5 | 4.9 |
| L-T-O | 0.3 | 0.4 | 1.2 | 0.0 | 0.0 | 0.1 | 13.9 | 0.6 | 29.5 | 0.4 | 47.9 | 5.6 |
| L-T-E | 0.4 | 0.5 | 0.6 | 0.0 | 0.1 | 0.1 | 0.9 | 0.3 | 15.3 | 0.6 | 10.7 | 70.7 |

**Table 9.** The multiclass confusion matrix for VGG16 showing the image classification accuracies (%) during inference on the ExoNet database. The columns and rows are the predicted and labelled classes, respectively.

|  | D-T-L | W-S | W-T-O | I-S | I-T-W | I-T-L | L-S | L-T-D | L-T-W | L-T-I | L-T-O | L-T-E |
| --- | --- | --- | --- | --- | --- | --- | --- | --- | --- | --- | --- | --- |
| D-T-L | 75.5 | 0.6 | 0.6 | 0.0 | 0.0 | 0.2 | 1.1 | 5.3 | 9.9 | 0.0 | 3.5 | 3.3 |
| W-S | 0.2 | 66.1 | 8.0 | 0.0 | 0.1 | 0.2 | 0.2 | 0.1 | 22.3 | 0.1 | 1.7 | 1.0 |
| W-T-O | 0.3 | 23.6 | 34.2 | 0.1 | 0.1 | 0.2 | 0.1 | 0.2 | 23.5 | 0.8 | 10.1 | 6.7 |
| I-S | 0.0 | 0.4 | 0.6 | 59.2 | 30.1 | 3.3 | 1.5 | 0.0 | 1.7 | 0.9 | 0.5 | 1.8 |
| I-T-W | 0.1 | 1.8 | 1.0 | 15.1 | 66.5 | 1.9 | 0.0 | 0.1 | 5.0 | 5.4 | 1.4 | 1.8 |
| I-T-L | 0.5 | 0.0 | 0.2 | 4.5 | 8.1 | 64.8 | 3.9 | 0.0 | 13.1 | 1.9 | 1.5 | 1.5 |
| L-S | 0.1 | 0.3 | 0.0 | 0.0 | 0.0 | 0.0 | 77.0 | 0.1 | 14.8 | 0.3 | 7.1 | 0.2 |
| L-T-D | 7.3 | 0.8 | 0.7 | 0.0 | 0.2 | 0.0 | 1.0 | 43.8 | 33.3 | 0.6 | 7.8 | 4.5 |
| L-T-W | 0.3 | 1.8 | 0.4 | 0.0 | 0.0 | 0.1 | 3.8 | 0.4 | 86.9 | 0.2 | 4.5 | 1.7 |
| L-T-I | 0.7 | 1.0 | 0.3 | 0.7 | 5.2 | 0.6 | 5.6 | 1.0 | 29.9 | 37.2 | 12.4 | 5.4 |
| L-T-O | 0.3 | 0.3 | 0.9 | 0.1 | 0.0 | 0.0 | 14.7 | 0.4 | 35.2 | 0.5 | 41.7 | 5.8 |
| L-T-E | 0.6 | 0.6 | 0.5 | 0.0 | 0.0 | 0.0 | 1.0 | 0.3 | 19.6 | 0.3 | 10.0 | 67.0 |

**Table 10.** The multiclass confusion matrix for VGG19 showing the image classification accuracies (%) during inference on the ExoNet database. The columns and rows are the predicted and labelled classes, respectively.

|  | D-T-L | W-S | W-T-O | I-S | I-T-W | I-T-L | L-S | L-T-D | L-T-W | L-T-I | L-T-O | L-T-E |
| --- | --- | --- | --- | --- | --- | --- | --- | --- | --- | --- | --- | --- |
| D-T-L | 73.9 | 0.5 | 0.4 | 0.0 | 0.1 | 0.1 | 1.2 | 4.4 | 12.0 | 0.2 | 3.6 | 3.7 |
| W-S | 0.4 | 65.0 | 7.3 | 0.0 | 0.3 | 0.2 | 1.0 | 0.3 | 21.9 | 0.3 | 2.0 | 1.1 |
| W-T-O | 1.1 | 24.5 | 31.0 | 0.1 | 0.3 | 0.7 | 0.1 | 1.4 | 24.2 | 0.8 | 8.7 | 7.3 |
| I-S | 0.0 | 0.5 | 0.0 | 73.0 | 19.7 | 3.1 | 0.5 | 0.0 | 1.4 | 0.8 | 0.5 | 0.5 |
| I-T-W | 0.1 | 1.1 | 1.1 | 20.7 | 61.7 | 2.4 | 0.1 | 0.1 | 5.1 | 5.3 | 1.1 | 1.2 |
| I-T-L | 0.3 | 1.2 | 0.2 | 5.2 | 5.7 | 66.3 | 4.5 | 0.0 | 12.0 | 1.9 | 1.2 | 1.5 |
| L-S | 0.0 | 0.5 | 0.0 | 0.0 | 0.0 | 0.1 | 76.8 | 0.0 | 13.5 | 0.4 | 8.5 | 0.2 |
| L-T-D | 6.4 | 0.5 | 0.8 | 0.0 | 0.1 | 0.0 | 1.6 | 41.2 | 35.9 | 0.4 | 8.2 | 4.9 |
| L-T-W | 0.4 | 1.8 | 0.4 | 0.0 | 0.0 | 0.1 | 4.0 | 0.4 | 86.1 | 0.2 | 4.7 | 1.8 |
| L-T-I | 0.4 | 1.6 | 0.2 | 0.5 | 5.1 | 0.5 | 5.9 | 0.3 | 28.6 | 37.6 | 13.5 | 5.8 |
| L-T-O | 0.5 | 0.3 | 0.9 | 0.0 | 0.0 | 0.1 | 15.6 | 0.5 | 34.5 | 0.3 | 41.5 | 5.7 |
| L-T-E | 0.4 | 0.7 | 0.4 | 0.0 | 0.0 | 0.1 | 1.0 | 0.3 | 21.2 | 0.5 | 10.9 | 64.5 |

**Table 11.** The multiclass confusion matrix for Xception showing the image classification accuracies (%) during inference on the ExoNet database. The columns and rows are the predicted and labelled classes, respectively.

|  | D-T-L | W-S | W-T-O | I-S | I-T-W | I-T-L | L-S | L-T-D | L-T-W | L-T-I | L-T-O | L-T-E |
| --- | --- | --- | --- | --- | --- | --- | --- | --- | --- | --- | --- | --- |
| D-T-L | 78.4 | 0.8 | 0.4 | 0.0 | 0.1 | 0.2 | 0.9 | 3.1 | 10.2 | 0.2 | 3.1 | 2.7 |
| W-S | 0.3 | 68.1 | 10.7 | 0.0 | 0.1 | 0.0 | 0.0 | 0.2 | 17.7 | 0.3 | 1.2 | 1.2 |
| W-T-O | 0.3 | 23.1 | 39.0 | 0.1 | 0.5 | 0.2 | 0.1 | 0.6 | 18.6 | 0.7 | 7.6 | 9.3 |
| I-S | 0.1 | 0.9 | 0.3 | 57.9 | 31.6 | 3.7 | 0.5 | 0.0 | 1.9 | 1.0 | 1.3 | 0.8 |
| I-T-W | 0.1 | 2.2 | 1.0 | 14.9 | 67.7 | 2.1 | 0.0 | 0.5 | 4.2 | 5.3 | 1.2 | 0.8 |
| I-T-L | 0.2 | 0.5 | 1.2 | 2.7 | 5.6 | 73.4 | 3.2 | 0.3 | 8.6 | 1.7 | 1.9 | 0.8 |
| L-S | 0.1 | 0.2 | 0.0 | 0.0 | 0.0 | 0.2 | 71.7 | 0.1 | 13.9 | 0.4 | 12.7 | 0.6 |
| L-T-D | 6.7 | 0.9 | 0.6 | 0.0 | 0.1 | 0.0 | 0.7 | 50.3 | 28.1 | 2.8 | 6.7 | 3.0 |
| L-T-W | 0.4 | 1.8 | 0.5 | 0.0 | 0.0 | 0.1 | 3.5 | 0.5 | 83.1 | 0.5 | 6.9 | 2.6 |
| L-T-I | 0.2 | 1.8 | 0.4 | 0.4 | 3.7 | 0.5 | 4.0 | 1.6 | 22.9 | 46.2 | 13.2 | 5.0 |
| L-T-O | 0.6 | 0.3 | 1.4 | 0.0 | 0.0 | 0.1 | 12.8 | 0.5 | 28.5 | 0.7 | 48.7 | 6.4 |
| L-T-E | 0.3 | 0.6 | 0.5 | 0.0 | 0.1 | 0.1 | 0.8 | 0.4 | 14.7 | 0.6 | 11.8 | 70.1 |

**Table 12.** The multiclass confusion matrix for ResNet50 showing the image classification accuracies (%) during inference on the ExoNet database. The columns and rows are the predicted and labelled classes, respectively.

|  | D-T-L | W-S | W-T-O | I-S | I-T-W | I-T-L | L-S | L-T-D | L-T-W | L-T-I | L-T-O | L-T-E |
| --- | --- | --- | --- | --- | --- | --- | --- | --- | --- | --- | --- | --- |
| D-T-L | 70.8 | 0.4 | 0.3 | 0.0 | 0.0 | 0.2 | 1.2 | 13.1 | 9.1 | 0.2 | 2.9 | 1.8 |
| W-S | 0.0 | 63.4 | 9.1 | 0.0 | 0.2 | 0.2 | 0.0 | 8.6 | 15.7 | 0.2 | 1.8 | 0.8 |
| W-T-O | 0.4 | 22.3 | 38.5 | 0.0 | 0.7 | 0.1 | 0.3 | 3.4 | 18.0 | 0.4 | 8.2 | 7.7 |
| I-S | 0.1 | 1.2 | 0.3 | 67.9 | 23.8 | 2.3 | 0.8 | 0.0 | 1.8 | 0.6 | 0.9 | 0.4 |
| I-T-W | 0.0 | 1.6 | 0.5 | 17.7 | 65.7 | 2.5 | 0.0 | 1.2 | 2.6 | 5.9 | 0.9 | 1.3 |
| I-T-L | 0.8 | 1.0 | 0.3 | 2.9 | 4.7 | 72.6 | 4.2 | 1.0 | 9.4 | 1.7 | 0.5 | 0.8 |
| L-S | 0.1 | 0.2 | 0.0 | 0.0 | 0.0 | 0.1 | 77.6 | 0.6 | 12.6 | 0.4 | 8.1 | 0.4 |
| L-T-D | 5.7 | 0.3 | 0.9 | 0.0 | 0.0 | 0.0 | 0.7 | 57.5 | 24.2 | 1.7 | 5.1 | 3.9 |
| L-T-W | 0.4 | 1.6 | 0.5 | 0.0 | 0.0 | 0.1 | 3.1 | 5.2 | 80.4 | 0.5 | 5.7 | 2.5 |
| L-T-I | 0.6 | 2.0 | 0.1 | 0.3 | 4.1 | 0.3 | 4.3 | 3.6 | 20.7 | 48.7 | 11.0 | 4.3 |
| L-T-O | 0.2 | 0.4 | 0.9 | 0.0 | 0.0 | 0.0 | 13.2 | 1.7 | 28.5 | 0.8 | 48.2 | 5.9 |
| L-T-E | 0.6 | 0.7 | 0.5 | 0.0 | 0.0 | 0.1 | 0.8 | 1.4 | 14.8 | 0.8 | 10.1 | 70.3 |

**Table 13.** The multiclass confusion matrix for ResNet101 showing the image classification accuracies (%) during inference on the ExoNet database. The columns and rows are the predicted and labelled classes, respectively.

|  | D-T-L | W-S | W-T-O | I-S | I-T-W | I-T-L | L-S | L-T-D | L-T-W | L-T-I | L-T-O | L-T-E |
| --- | --- | --- | --- | --- | --- | --- | --- | --- | --- | --- | --- | --- |
| D-T-L | 77.7 | 0.5 | 0.2 | 0.0 | 0.1 | 0.3 | 1.1 | 4.5 | 9.7 | 0.1 | 3.5 | 2.3 |
| W-S | 0.4 | 66.6 | 10.1 | 0.0 | 0.2 | 0.2 | 0.1 | 0.3 | 18.3 | 0.1 | 2.2 | 1.5 |
| W-T-O | 0.3 | 24.5 | 41.8 | 0.0 | 0.4 | 0.1 | 0.0 | 0.4 | 15.6 | 0.6 | 7.5 | 9.0 |
| I-S | 0.1 | 0.5 | 0.1 | 63.1 | 28.7 | 2.2 | 0.5 | 0.0 | 1.5 | 1.3 | 1.3 | 0.6 |
| I-T-W | 0.1 | 1.8 | 0.8 | 20.1 | 63.2 | 3.5 | 0.0 | 0.2 | 2.4 | 5.1 | 0.6 | 2.2 |
| I-T-L | 0.5 | 1.0 | 0.2 | 3.5 | 3.9 | 58.2 | 2.7 | 0.2 | 10.6 | 1.9 | 2.0 | 15.3 |
| L-S | 0.1 | 0.4 | 0.0 | 0.0 | 0.0 | 0.1 | 75.0 | 0.1 | 12.8 | 0.6 | 8.9 | 2.0 |
| L-T-D | 7.3 | 0.7 | 0.6 | 0.0 | 0.1 | 0.0 | 1.0 | 53.1 | 26.2 | 1.8 | 5.8 | 3.5 |
| L-T-W | 0.3 | 1.7 | 0.5 | 0.0 | 0.0 | 0.1 | 3.5 | 0.6 | 81.5 | 0.4 | 6.5 | 4.9 |
| L-T-I | 0.2 | 1.8 | 0.2 | 0.4 | 3.4 | 0.2 | 4.1 | 1.6 | 21.9 | 47.9 | 12.1 | 6.2 |
| L-T-O | 0.5 | 0.4 | 1.0 | 0.0 | 0.0 | 0.1 | 12.5 | 0.6 | 27.8 | 0.7 | 48.6 | 8.0 |
| L-T-E | 0.8 | 0.6 | 0.6 | 0.0 | 0.1 | 0.1 | 0.7 | 0.3 | 13.6 | 0.6 | 11.1 | 71.5 |

**Table 14.** The multiclass confusion matrix for ResNet152 showing the image classification accuracies (%) during inference on the ExoNet database. The columns and rows are the predicted and labelled classes, respectively.

|  | D-T-L | W-S | W-T-O | I-S | I-T-W | I-T-L | L-S | L-T-D | L-T-W | L-T-I | L-T-O | L-T-E |
| --- | --- | --- | --- | --- | --- | --- | --- | --- | --- | --- | --- | --- |
| D-T-L | 77.1 | 0.6 | 0.6 | 0.0 | 0.1 | 0.1 | 1.0 | 4.2 | 10.4 | 0.2 | 3.4 | 2.3 |
| W-S | 0.6 | 60.5 | 10.9 | 0.0 | 0.1 | 0.3 | 0.0 | 0.7 | 25.0 | 0.3 | 1.1 | 0.6 |
| W-T-O | 0.4 | 20.3 | 42.3 | 0.0 | 0.2 | 0.0 | 0.0 | 0.4 | 23.7 | 0.6 | 8.2 | 4.1 |
| I-S | 0.1 | 0.9 | 0.1 | 62.5 | 27.9 | 3.3 | 1.0 | 0.0 | 1.8 | 1.7 | 0.5 | 0.1 |
| I-T-W | 0.1 | 1.8 | 0.8 | 18.6 | 64.8 | 2.6 | 0.3 | 0.5 | 2.9 | 5.4 | 1.2 | 1.2 |
| I-T-L | 0.2 | 1.0 | 0.3 | 2.5 | 5.2 | 72.2 | 3.9 | 0.2 | 9.9 | 1.9 | 1.7 | 1.0 |
| L-S | 0.1 | 0.3 | 0.0 | 0.0 | 0.0 | 0.1 | 78.5 | 0.3 | 12.7 | 0.3 | 7.4 | 0.4 |
| L-T-D | 5.8 | 0.5 | 0.4 | 0.0 | 0.2 | 0.1 | 0.9 | 54.5 | 26.2 | 1.8 | 6.4 | 3.2 |
| L-T-W | 0.3 | 1.5 | 0.6 | 0.0 | 0.0 | 0.1 | 3.8 | 0.5 | 85.4 | 0.5 | 5.0 | 2.3 |
| L-T-I | 0.1 | 0.9 | 0.1 | 0.4 | 3.8 | 0.7 | 4.7 | 1.5 | 23.5 | 46.7 | 12.6 | 5.1 |
| L-T-O | 0.4 | 0.4 | 0.9 | 0.0 | 0.1 | 0.0 | 14.0 | 0.8 | 29.9 | 0.5 | 46.9 | 6.0 |
| L-T-E | 0.4 | 0.5 | 0.4 | 0.0 | 0.0 | 0.1 | 0.7 | 0.7 | 15.6 | 0.7 | 9.9 | 70.9 |

**Table 15.** The multiclass confusion matrix for DenseNet121 showing the image classification accuracies (%) during inference on the ExoNet database. The columns and rows are the predicted and labelled classes, respectively.

|  | D-T-L | W-S | W-T-O | I-S | I-T-W | I-T-L | L-S | L-T-D | L-T-W | L-T-I | L-T-O | L-T-E |
| --- | --- | --- | --- | --- | --- | --- | --- | --- | --- | --- | --- | --- |
| D-T-L | 79.2 | 0.5 | 0.1 | 0.0 | 0.2 | 0.3 | 1.1 | 4.1 | 9.0 | 0.3 | 3.1 | 2.2 |
| W-S | 0.2 | 66.4 | 8.8 | 0.0 | 0.3 | 0.1 | 0.0 | 0.4 | 21.5 | 0.3 | 1.4 | 0.6 |
| W-T-O | 0.5 | 21.1 | 35.2 | 0.0 | 0.4 | 0.0 | 1.0 | 0.2 | 24.1 | 0.8 | 11.1 | 5.7 |
| I-S | 0.1 | 0.3 | 0.6 | 60.8 | 29.2 | 2.3 | 0.5 | 0.0 | 2.3 | 0.9 | 1.8 | 1.2 |
| I-T-W | 0.2 | 1.8 | 0.4 | 20.3 | 64.9 | 3.1 | 0.0 | 0.1 | 2.8 | 4.6 | 0.8 | 1.1 |
| I-T-L | 0.2 | 0.7 | 0.3 | 3.0 | 4.9 | 74.6 | 3.2 | 0.3 | 8.6 | 1.9 | 1.3 | 1.0 |
| L-S | 0.1 | 0.1 | 0.0 | 0.0 | 0.0 | 0.2 | 78.3 | 0.3 | 11.9 | 0.2 | 8.3 | 0.7 |
| L-T-D | 6.0 | 0.7 | 0.4 | 0.0 | 0.2 | 0.1 | 0.8 | 53.0 | 27.2 | 2.4 | 5.7 | 3.4 |
| L-T-W | 0.4 | 1.6 | 0.5 | 0.0 | 0.0 | 0.1 | 3.9 | 0.5 | 84.4 | 0.5 | 5.8 | 2.2 |
| L-T-I | 0.1 | 1.2 | 0.3 | 0.7 | 4.1 | 0.7 | 4.6 | 1.1 | 23.6 | 48.8 | 10.7 | 4.1 |
| L-T-O | 0.5 | 0.4 | 0.8 | 0.0 | 0.0 | 0.1 | 13.8 | 0.6 | 29.3 | 0.6 | 48.6 | 5.2 |
| L-T-E | 0.7 | 0.6 | 0.4 | 0.0 | 0.1 | 0.2 | 0.7 | 0.4 | 15.8 | 0.8 | 10.7 | 69.6 |

**Table 16.** The multiclass confusion matrix for DenseNet169 showing the image classification accuracies (%) during inference on the ExoNet database. The columns and rows are the predicted and labelled classes, respectively.

|  | D-T-L | W-S | W-T-O | I-S | I-T-W | I-T-L | L-S | L-T-D | L-T-W | L-T-I | L-T-O | L-T-E |
| --- | --- | --- | --- | --- | --- | --- | --- | --- | --- | --- | --- | --- |
| D-T-L | 75.0 | 0.5 | 0.1 | 0.0 | 0.1 | 0.1 | 0.8 | 4.0 | 13.7 | 0.1 | 3.3 | 2.2 |
| W-S | 0.3 | 68.9 | 8.8 | 0.0 | 0.1 | 0.2 | 0.0 | 0.2 | 18.5 | 0.2 | 1.8 | 1.1 |
| W-T-O | 0.3 | 24.1 | 36.9 | 0.0 | 0.2 | 0.1 | 0.6 | 0.4 | 19.6 | 0.4 | 7.1 | 10.4 |
| I-S | 0.0 | 0.8 | 0.1 | 58.5 | 32.8 | 2.4 | 0.4 | 0.1 | 2.3 | 1.4 | 1.0 | 0.1 |
| I-T-W | 0.2 | 2.0 | 0.9 | 17.8 | 64.1 | 3.9 | 0.0 | 0.1 | 5.5 | 3.9 | 1.0 | 0.8 |
| I-T-L | 0.3 | 0.5 | 0.0 | 2.9 | 5.9 | 72.9 | 3.0 | 0.2 | 9.8 | 1.9 | 1.9 | 0.8 |
| L-S | 0.2 | 0.3 | 0.0 | 0.0 | 0.0 | 0.2 | 78.1 | 0.2 | 12.6 | 0.3 | 7.6 | 0.5 |
| L-T-D | 7.0 | 0.9 | 0.6 | 0.0 | 0.2 | 0.1 | 0.8 | 50.8 | 28.5 | 2.0 | 5.7 | 3.5 |
| L-T-W | 0.4 | 1.8 | 0.5 | 0.0 | 0.1 | 0.1 | 3.6 | 0.6 | 84.3 | 0.4 | 5.4 | 2.9 |
| L-T-I | 0.2 | 1.7 | 0.1 | 0.3 | 4.8 | 0.2 | 3.6 | 1.8 | 24.3 | 45.9 | 12.6 | 4.4 |
| L-T-O | 0.5 | 0.3 | 0.9 | 0.0 | 0.0 | 0.1 | 13.6 | 0.7 | 30.1 | 0.5 | 46.9 | 6.3 |
| L-T-E | 0.6 | 0.6 | 0.5 | 0.0 | 0.0 | 0.1 | 0.7 | 0.5 | 18.0 | 0.6 | 10.2 | 68.2 |

**Table 17.** The multiclass confusion matrix for DenseNet201 showing the image classification accuracies (%) during inference on the ExoNet database. The columns and rows are the predicted and labelled classes, respectively.

|  | D-T-L | W-S | W-T-O | I-S | I-T-W | I-T-L | L-S | L-T-D | L-T-W | L-T-I | L-T-O | L-T-E |
| --- | --- | --- | --- | --- | --- | --- | --- | --- | --- | --- | --- | --- |
| D-T-L | 75.6 | 0.9 | 0.2 | 0.0 | 0.0 | 0.4 | 0.7 | 5.1 | 10.0 | 0.1 | 4.0 | 2.8 |
| W-S | 0.3 | 67.9 | 12.1 | 0.0 | 0.2 | 0.1 | 0.0 | 0.3 | 16.2 | 0.6 | 1.7 | 0.5 |
| W-T-O | 2.4 | 23.6 | 37.8 | 0.0 | 0.4 | 0.2 | 0.0 | 1.5 | 19.9 | 0.4 | 8.3 | 5.5 |
| I-S | 0.1 | 0.0 | 0.3 | 59.4 | 33.7 | 2.7 | 0.1 | 0.1 | 1.5 | 1.2 | 0.9 | 0.0 |
| I-T-W | 0.2 | 1.9 | 1.0 | 14.2 | 68.9 | 3.0 | 0.1 | 0.1 | 2.8 | 5.6 | 1.2 | 1.0 |
| I-T-L | 0.2 | 0.5 | 0.2 | 4.0 | 5.7 | 72.2 | 2.2 | 0.2 | 10.3 | 2.0 | 1.3 | 1.2 |
| L-S | 0.0 | 0.2 | 0.0 | 0.0 | 0.0 | 0.1 | 76.0 | 0.2 | 13.4 | 0.6 | 8.9 | 0.6 |
| L-T-D | 6.4 | 0.7 | 0.8 | 0.0 | 0.2 | 0.1 | 1.2 | 51.4 | 27.0 | 2.7 | 5.0 | 4.6 |
| L-T-W | 0.4 | 1.8 | 0.9 | 0.0 | 0.0 | 0.1 | 3.5 | 0.6 | 82.3 | 0.5 | 7.0 | 2.8 |
| L-T-I | 0.7 | 1.5 | 0.4 | 0.5 | 4.5 | 0.4 | 5.1 | 1.3 | 23.9 | 47.2 | 10.8 | 3.7 |
| L-T-O | 0.5 | 0.4 | 1.4 | 0.0 | 0.0 | 0.1 | 12.6 | 0.8 | 29.6 | 0.8 | 47.4 | 6.4 |
| L-T-E | 0.8 | 0.6 | 0.4 | 0.0 | 0.1 | 0.1 | 0.9 | 0.6 | 13.5 | 0.7 | 12.0 | 70.4 |

## 5 Discussion

In this study, we developed an advanced environment classification system for robotic leg prostheses and exoskeletons using computer vision and deep learning. Taking inspiration from the human vision-locomotor control system, environment sensing and classification could improve the automated high-level controller by reconstructing the oncoming walking environments prior to physical interaction, therein allowing for more accurate and real-time transitions between different locomotion modes. However, small-scale and private image datasets have impeded the development and comparison of environment classification algorithms from different researchers, respectively. We therefore developed ExoNet - the largest open-source dataset of wearable camera images of walking environments. Unparalleled in scale and diversity, ExoNet contains ~5.6 million images of indoor and outdoor real-world walking environments, of which ~923,000 images were annotated using a 12-class hierarchical labelling architecture. We then trained and tested over a dozen state-of-the-art deep convolutional neural networks on the ExoNet database for large-scale image classification of the walking environments, including: EfficientNetB0, InceptionV3, MobileNet, MobileNetV2, VGG16, VGG19, Xception, ResNet50, ResNet101, ResNet152, DenseNet121, DenseNet169, and DenseNet201. This research provides a benchmark and reference for future comparisons. Although we designed our environment classification system for robotic prostheses and exoskeletons, applications could extend to humanoids, autonomous legged robots, powered wheelchairs, and assistive devices for persons with visual impairments.

The use of deep convolutional neural networks was made possible by the ExoNet database. In addition to being open-source, the large scale and diversity of ExoNet significantly distinguishes itself from previous research (see Table 1). The ExoNet database contains ~923,000 labelled images. In comparison, the previous largest dataset, developed by Varol’s research group (Massalin et al., 2018), contained ~402,000 images. Whereas previous datasets included fewer than six environment classes, the most common being level-ground terrain and incline and decline stairs, the ExoNet database contains a 12-class hierarchical labelling architecture; these differences have important practical implications since deep learning requires significant and diverse training data to prevent overfitting and promote generalization (LeCun et al., 2015). Furthermore, our image quality (1280×720) is considerably higher than previous datasets (e.g., 224×224 and 320×240). Lower resolution images have shown to decrease the classification accuracy of walking environments (Da Silva et al, 2020; Novo-Torres et al., 2019). Although higher resolution images can increase the onboard computational and memory storage requirements, using efficient CNN architectures with fewer operations like EfficientNetB0 (Tan and Le, 2019) allow for processing larger images for relatively similar computational cost. As robotic leg prostheses and exoskeletons begin to transition out of research laboratories and into real-world environments, large-scale and challenging datasets like ExoNet are needed to support the development of next-generation image classification algorithms for environment-adaptive locomotor control.

A potential limitation of the ExoNet database is the 2D nature of the environment information. Many researchers have likewise used a wearable RGB camera for passive environment sensing (Da Silva et al., 2020; Diaz et al., 2018; Khademi and Simon, 2019; Krausz and Hargrove, 2015; Laschowski et al., 2019b; 2020b; Novo-Torres et al., 2019; Zhong et al., 2020). Although multi-camera systems could be used to capture 3D information (i.e., comparable to how the human visual system uses triangulation for depth perception) (Patla, 1997), each pixel in an RGB image contains only light intensity information. Other researchers have used depth cameras to explicitly capture images containing both light intensity information and distance measurements (Hu et al., 2018; Krausz et al., 2015; 2019; Krausz and Hargrove, 2021; Massalin et al., 2018; Varol and Massalin, 2016; Zhang et al., 2019b; 2019c; 2019d, 2020). These range imaging systems work by emitting infrared light and measuring the light time-of-flight between the camera and oncoming walking environment to calculate distance. Depth sensing can uniquely extract environmental features like step height and width, which can improve the mid-level control of robotic prostheses and exoskeletons (e.g., increasing actuator joint torques to assist with steeper stairs).

Despite these benefits, depth measurement accuracy typically degrades in outdoor lighting conditions (e.g., sunlight) and with increasing distance (Krausz and Hargrove, 2019; Zhang et al., 2019a). Consequently, most environment recognition systems using depth cameras have been tested in controlled indoor environments and/or have had limited capture volumes (i.e., 1-2 m of maximum range imaging) (Hu et al., 2018; Krausz et al., 2015; 2019; Massalin et al., 2018; Varol and Massalin, 2016, Zhang et al., 2020). These systems usually require an onboard accelerometer or IMU to transform the 3D environment information from the camera coordinate system into global coordinates (Krausz et al., 2015; Krausz and Hargrove, 2021; Zhang et al., 2019b; 2019c; 2019d; 2020). The application of depth cameras for active environment sensing would also require robotic prostheses and exoskeletons to have onboard microcontrollers with high computing power and low power consumption. The current embedded systems would need significant modifications to support the real-time processing and classification of depth images given their added complexity. These practical limitations motivated our decision to use RGB images for the environment classification.

The EfficientNetB0 network (Tan and Le, 2019) achieved the highest image classification accuracy on the ExoNet database during inference (i.e., 73.2% accuracy). However, for environment-adaptive control of robotic leg prostheses and exoskeletons, higher classification accuracies are desired since even rare misclassifications could cause loss-of-balance and injury. Although we used state-of-the-art deep convolutional neural networks, the architectures included only feedforward connections, therein classifying the walking environment frame-by-frame without knowing the preceding classification decisions. Moving forward, sequential information over time could be used to improve the classification accuracy and robustness, especially during steady-state environments. This computer vision technique is analogous to how light-sensitive receptors in the human eye capture dynamic images to control locomotion (i.e., known as optical flow) (Patla, 1997). Sequential data could be classified using majority voting (Wang et al., 2013; Massalin et al., 2018; Varol and Massalin, 2016) or deep learning models like Transformers (Vaswani et al., 2017) or recurrent neural networks (RNNs) (Zhang et al., 2019b). Majority voting works by storing sequential decisions in a vector and generates a classification prediction based on the majority of decisions.

Recurrent neural networks process sequential data while maintaining an internal hidden state vector that implicitly contains temporal information. Training RNNs can be challenging though, due to exploding and vanishing gradients. Although these neural networks were designed to learn long-term dependencies, research has shown that they struggle with storing sequential information over long periods (LeCun et al., 2015). To mitigate this issue, RNNs can be supplemented with an explicit memory module like a neural Turning machine or long short-term memory (LSTM) network, therein allowing for uninterrupted backpropagated gradients. Fu’s research group (Zhang et al., 2019b) explored using sequential data for environment classification. Sequential decisions from a baseline CNN were fused and classified using a recurrent neural network, LSTM network, majority voting, and a hidden Markov model (HMM). The baseline neural network achieved 92.8% image classification accuracy across five environment classes. Supplementing the baseline CNN with an RNN, LSTM network, majority voting, and HMM resulted in 96.5%, 96.4%, 95%, and 96.8% classification accuracies, respectively (Zhang et al., 2019b). Although using sequential data can improve the classification accuracy of walking environments, these decisions require longer computation times and thus could impede real-time locomotor control.

The development of deep convolutional neural networks has traditionally focused on improving classification accuracy, often leading to more accurate yet inefficient algorithms with greater computational and memory storage requirements (Canziani et al., 2016). These design features can be especially problematic for deployment on mobile and embedded systems, which inherently have limited operating resources. Despite the advances in computing platforms like graphics processing units (GPUs), the current embedded systems in robotic leg prostheses and exoskeletons would struggle to support the architectural and computational complexities typically associated with deep learning for computer vision. Accordingly, we quantitatively evaluated and compared the benchmarked CNN architectures and their environment classification predictions on the ExoNet database using an operational metric called “NetScore” (Wong, 2018), which balances the image classification accuracy with the computational and memory storage requirements (i.e., important for onboard real-time inference). Although the ResNet152 network achieved one of the highest image classification accuracies on the ExoNet database (i.e., 71.6% accuracy), it received the lowest NetScore (i.e., Ω = 46) due to the disproportionally large number of parameters (i.e., containing more parameters than any other CNN architecture we tested). Interestingly, the EfficientNetB0 network did not receive the highest NetScore (i.e., Ω = 72.6) despite achieving the highest classification accuracy and the architecture having been optimally designed using a neural architecture search to maximize the accuracy while minimizing the number of computing operations (Tan and Le, 2019).

The MobileNetV2 network developed by Google (Sander et al., 2019), which uses depthwise separable convolutions, received the highest NetScore (i.e., Ω = 76.2), therein demonstrating the best balance between the classification accuracy (i.e., 72.9% accuracy) and the architectural and computational complexities. The authors previously demonstrated the ability of their CNN architecture to perform onboard real-time inference on a mobile device (i.e., ~75 ms per image on a CPU-powered Google Pixel 1 smartphone) (Sander et al., 2019). However, our classification system could theoretically generate even faster runtimes since 1) the smartphone we used (i.e., the iPhone XS Max) has an onboard GPU, and 2) we reduced the size of the final densely connected layer of the MobileNetV2 architecture from 1,000 outputs (i.e., as originally used for ImageNet) to 12 outputs (i.e., for the ExoNet database). Compared to traditional CPUs, GPUs have many more core processors, which permit faster and more efficient CNN computations through parallel computing (LeCun et al., 2015). Moving forward, we recommend using the existing CPU embedded systems in robotic prostheses and exoskeletons for performing the high-level locomotion mode recognition based on neuromuscular-mechanical data, which is less computationally expensive, and a supplementary GPU computing platform for the image-based environment classification; these recommendations concur with those recently proposed by Huang and colleagues (Da Silva et al, 2020).

Lastly, given that the environmental context does not explicitly represent the user’s locomotor intent, although it constrains the movement possibilities, data from computer vision should supplement, rather than replace, the automated locomotion mode control decisions based on mechanical, inertial, and/or neuromuscular signals. Images from our wearable smartphone camera could be fused with its onboard IMU measurements to improve the high-level control performance. For example, when an exoskeleton or prosthesis user wants to sit down, the acceleration data would indicate stand-to-sit rather than level-ground walking, despite the level-ground terrain being accuracy detected within the field-of-view (bottom right image in Figure 7). Inspired by previous work (Da Silva et al., 2020; Diaz et al., 2018; Khademi and Simon, 2019; Wang et al., 2013; Zhang et al., 2011), our smartphone IMU measurements could also help minimize the onboard computational and memory storage requirements via sampling rate control (i.e., providing an automatic triggering mechanism for the image capture). Whereas faster walking speeds can benefit from higher sampling rates for continuous classification, standing still does not necessarily require environment information and thus the smartphone camera could be powered down or the sampling rate decreased, therein conserving the onboard operating resources. However, relatively few researchers have fused environment data with mechanical and/or inertial measurements for automated locomotion mode recognition (Du et al., 2012; Huang et al., 2011a; Krausz et al., 2019; Krausz and Hargrove, 2021; Liu et al., 2016; Wang et al., 2013; Zhang et al., 2011) and only one study (Zhang et al., 2020) has used such information for online environment-adaptive control of a robotic prosthesis during walking (i.e., stepping over an obstacle). These limitations in systems integration offer exciting challenges and opportunities for future research.

## 7 Author Contributions

BL contributed to the study design, literature review, data collection, image labelling, CNN training and testing, analyses of the results, and manuscript writing. WM contributed to the study design, code development, CNN training and testing, analyses of the results, and manuscript writing. AW and JM contributed to the study design, analyses of the results, and manuscript writing. All authors read and approved the final manuscript.

## 9 Acknowledgments

We acknowledge the TensorFlow Research Cloud program by Google and the NVIDIA GPU Grant program for providing the computing hardware. We also thank Helen Huang and Edgar Lobaton (North Carolina State University, USA); Levi Hargrove (Northwestern University, USA); Huseyin Atakan Varol (Nazarbayev University, Kazakhstan); Dan Simon (Cleveland State University, USA); Dario Villarreal (Southern Methodist University, USA); and Chenglong Fu (Southern University of Science and Technology, China) for their correspondences.

## 10 Data Availability Statement

The image dataset generated for this research study, which we named ExoNet, was uploaded to the IEEE DataPort repository and is publicly available for download at https://ieee-dataport.org/open-access/exonet-database-wearable-camera-images-human-locomotion-environments

## References

Canziani, A., Paszke, A., and Culurciello, E. (2016). An analysis of deep neural network models for practical applications. arXiv [Preprint]. arXiv:1605.07678.

Chollet, F. (2016). Xception: Deep learning with depthwise separable convolutions. arXiv [Preprint]. arXiv:1610.02357.

Da Silva, R. L., Starliper, N., Zhong, B., Huang, H. H., and Lobaton, E. (2020). Evaluation of embedded platforms for lower limb prosthesis with visual sensing capabilities. arXiv [Preprint]. arXiv:2006.15224.

Deng, J., Dong, W., Socher, R., Li, L. J., Li, K., and Fei-Fei, L. (2009). “ImageNet: A large-scale hierarchical image database,” in IEEE Conference on Computer Vision and Pattern Recognition (CVPR) (Miami, FL: IEEE), 248–255. doi: 10.1109/CVPR.2009.5206848.

Diaz, J. P., Da Silva, R. L., Zhong, B., Huang, H. H., and Lobaton, E. (2018). “Visual terrain identification and surface inclination estimation for improving human locomotion with a lower-limb prosthetic,” in Annual International Conference of the IEEE Engineering in Medicine and Biology Society (EMBC) (Honolulu, HI: IEEE), 1817–1820. doi: 10.1109/EMBC.2018.8512614.

Ding, X., Zhang, X., Ma, N., Han, J., Ding, G., and Sun, J. (2021). RepVGG: Making VGG-style ConvNets great again. arXiv [Preprint]. arXiv:2101.03697.

Du, L., Zhang, F., Liu, M., and Huang, H. (2012). Toward design of an environment-aware adaptive locomotion-mode-recognition system. IEEE Transactions on Biomedical Engineering, 59, 2716–2725. doi: 10.1109/TBME.2012.2208641.

Grimmer, M., Riener, R., Walsh, C. J., and Seyfarth, A. (2019). Mobility related physical and functional losses due to aging and disease - a motivation for lower limb exoskeletons. Journal of Neuroengineering and Rehabilitation, 16, 2. doi: 10.1186/s12984-018-0458-8.

He, K., Zhang, X., Ren, S., and Sun, J. (2015). Deep residual learning for image recognition. arXiv [Preprint]. arXiv:1512.03385.

Howard, A. G., Zhu, M., Chen, B., Kalenichenko, D., Wang, W., Weyand, T., Andreetto, M., and Adam, H. (2017). MobileNets: Efficient convolutional neural networks for mobile vision applications. arXiv [Preprint]. arXiv:1704.04861.

Hu, B. H., Krausz, N. E., and Hargrove, L. J. (2018). “A novel method for bilateral gait segmentation using a single thigh-mounted depth sensor and IMU,” in IEEE International Conference on Biomedical Robotics and Biomechatronics (BIOROB) (Enschede: IEEE), 807–812. doi: 10.1109/BIOROB.2018.8487806.

Huang, H., Kuiken, T. A., and Lipschutz, R. D. (2009). A strategy for identifying locomotion modes using surface electromyography. IEEE Transactions on Biomedical Engineering, 56, 65–73. doi: 10.1109/TBME.2008.2003293.

Huang, H., Dou, Z., Zheng, F., and Nunnery, M. J. (2011a). “Improving the performance of a neural-machine interface for artificial legs using prior knowledge of walking environment,” in Annual International Conference of the IEEE Engineering in Medicine and Biology Society (EMBC) (Boston, MA: IEEE), 4255–4258. doi: 10.1109/IEMBS.2011.6091056.

Huang, H., Zhang, F., Hargrove, L. J., Dou, Z., Rogers, D. R., and Englehart, K. B. (2011b). Continuous locomotion-mode identification for prosthetic legs based on neuromuscular-mechanical fusion. IEEE Transactions on Biomedical Engineering, 58, 2867–2875. doi: 10.1109/TBME.2011.2161671.

Huang, G., Liu, Z., van der Maaten, L., and Weinberger, K. Q. (2017). “Densely connected convolutional networks,” in IEEE Conference on Computer Vision and Pattern Recognition (CVPR) (Honolulu, HI: IEEE). doi: 10.1109/CVPR.2017.243.

Karacan, K., Meyer, J. T., Bozma, H. I., Gassert, R., and Samur, E. (2020). “An environment recognition and parameterization system for shared-control of a powered lower-limb exoskeleton,” in IEEE RAS/EMBS International Conference for Biomedical Robotics and Biomechatronics (BioRob) (New York, NY: IEEE), doi: 10.1109/BioRob49111.2020.9224407.

Khademi, G., and Simon, D. (2019). “Convolutional neural networks for environmentally aware locomotion mode recognition of lower-limb amputees,” in ASME Dynamic Systems and Control Conference (DSCC) (Park City, UT: ASME), 11. doi: 10.1115/DSCC2019-9180.

Kingma, D. P., and Ba, J. (2015). Adam: A method for stochastic optimization. arXiv [Preprint]. arXiv:1412.6980.

Kleiner, B., Ziegenspeck, N., Stolyarov, R., Herr, H., Schneider, U., and Verl, A. (2018). “A radar-based terrain mapping approach for stair detection towards enhanced prosthetic foot control,” in IEEE International Conference on Biomedical Robotics and Biomechatronics (BIOROB) (Enschede: IEEE), 105–110. doi: 10.1109/BIOROB.2018.8487722.

Krausz, N. E., and Hargrove, L. J. (2015). “Recognition of ascending stairs from 2D images for control of powered lower limb prostheses,” in International IEEE/EMBS Conference on Neural Engineering (NER) (Montpellier: IEEE), 615–618. doi: 10.1109/NER.2015.7146698.

Krausz, N. E., Lenzi, T., and Hargrove, L. J. (2015). Depth sensing for improved control of lower limb prostheses. IEEE Transactions on Biomedical Engineering, 62, 2576–2587. doi: 10.1109/TBME.2015.2448457.

Krausz, N. E., and Hargrove, L. J. (2019). A survey of teleceptive sensing for wearable assistive robotic devices. Sensors, 19, 5238. doi: 10.3390/s19235238.

Krausz, N. E., Hu, B. H., and Hargrove, L. J. (2019). Subject- and environment-based sensor variability for wearable lower-limb assistive devices. Sensors, 19, 4887. doi: 10.3390/s19224887.

Krausz, N. E., and Hargrove, L. J. (2021). Sensor fusion of vision, kinetics and kinematics for forward prediction during walking with a transfemoral prosthesis. IEEE Transactions on Medical Robotics and Bionics. doi: 10.1109/TMRB.2021.3082206.

Krizhevsky, A., Sutskever, I., and Hinton, G. E. (2012). “ImageNet classification with deep convolutional neural networks,” in Advances in Neural Information Processing Systems Conference (NIPS) (Lake Tahoe, NV), 1097–1105. doi: 10.1145/3065386.

Laschowski, B., and Andrysek, J. (2018). “Electromechanical design of robotic transfemoral prostheses,” in ASME International Design Engineering Technical Conferences and Computers and Information in Engineering Conference (IDETC-CIE) (Quebec City: ASME), V05AT07A054. doi: 10.1115/DETC2018-85234.

Laschowski, B., McPhee, J., and Andrysek, J. (2019a). Lower-limb prostheses and exoskeletons with energy regeneration: Mechatronic design and optimization review. ASME Journal of Mechanisms and Robotics, 11, 040801. doi: 10.1115/1.4043460.

Laschowski, B., McNally, W., Wong, A., and McPhee, J. (2019b). “Preliminary design of an environment recognition system for controlling robotic lower-limb prostheses and exoskeletons,” in IEEE International Conference on Rehabilitation Robotics (ICORR) (Toronto: IEEE), 868–873. doi: 10.1109/ICORR.2019.8779540.

Laschowski, B., McNally, W., Wong, A., and McPhee, J. (2020a). “Comparative analysis of environment recognition systems for control of lower-limb exoskeletons and prostheses,” in IEEE International Conference on Biomedical Robotics and Biomechatronics (BIOROB) (New York City, NY: IEEE). doi: 10.1109/BioRob49111.2020.9224364.

Laschowski, B., McNally, W., Wong, A., and McPhee, J. (2020b). ExoNet database: Wearable camera images of human locomotion environments. Frontiers in Robotics and AI, 7, 562061. doi: 10.3389/frobt.2020.562061.

Laschowski, B., Razavian, R. S., and McPhee, J. (2021a). Simulation of stand-to-sit biomechanics for robotic exoskeletons and prostheses with energy regeneration. IEEE Transactions on Medical Robotics and Bionics, 3, 455–462. doi: 10.1109/TMRB.2021.3058323.

Laschowski, B., McNally, W., Wong, A., and McPhee, J. (2021b). Computer vision and deep learning for environment-adaptive control of robotic lower-limb exoskeletons. bioRxiv [Preprint]. doi: 10.1101/2021.04.02.438126.

Lebedev, M. A., and Nicolelis, M. A. L. (2017). Brain-machine interfaces: From basic science to neuroprosthetics and neurorehabilitation. Physiological Reviews, 97, 767–837. doi: 10.1152/physrev.00027.2016.

LeCun, Y., Bengio, Y., and Hinton, G. (2015). Deep learning. Nature, 521, 436–444. doi: 10.1038/nature14539.

Li, M., Zhong, B., Liu, Z., Lee, I. C., Fylstra, B. L., Lobaton, E., et al. (2019). “Gaze fixation comparisons between amputees and able-bodied individuals in approaching stairs and level-ground transitions: a pilot study,” in Annual International Conference of the IEEE Engineering in Medicine and Biology Society (EMBC) (Berlin: IEEE). doi: 10.1109/EMBC.2019.8857388.

Liu, M., Wang, D., and Huang, H. (2016). Development of an environment-aware locomotion mode recognition system for powered lower limb prostheses. IEEE Transactions on Neural Systems and Rehabilitation Engineering, 24, 434–443. doi: 10.1109/TNSRE.2015.2420539.

Loshchilov, I., and Hutter, F. (2016). SGDR: Stochastic gradient descent with warm restarts. arXiv [Preprint]. arXiv:1608.03983.

Massalin, Y., Abdrakhmanova, M., and Varol, H. A. (2018). User-independent intent recognition for lower limb prostheses using depth sensing. IEEE Transactions on Biomedical Engineering, 65, 1759–1770. doi: 10.1109/TBME.2017.2776157.

Nasr, A., Laschowski, B., and McPhee, J. (2021). “Myoelectric control of robotic leg prostheses and exoskeletons: A review” in ASME International Design Engineering Technical Conferences and Computers and Information in Engineering Conference (IDETC-CIE) (Virtual: ASME) Accepted.

Negrini, S., Mills, J. A., Arienti, C., Kiekens, C., and Cieza, A. (2021). “Rehabilitation research framework for patients with COVID-19” defined by Cochrane rehabilitation and the World Health Organization rehabilitation programme. Archives of Physical Medicine and Rehabilitation. doi: 10.1016/j.apmr.2021.02.018.

Novo-Torres, L., Ramirez-Paredes, J. P., and Villarreal, D. J. (2019). “Obstacle recognition using computer vision and convolutional neural networks for powered prosthetic leg applications,” in Annual International Conference of the IEEE Engineering in Medicine and Biology Society (EMBC) (Berlin: IEEE), 3360–3363. doi: 10.1109/EMBC.2019.8857420.

Patla, A. E. (1997). Understanding the roles of vision in the control of human locomotion. Gait & Posture, 5, 54–69. doi: 10.1016/S0966-6362(96)01109-5.

Rai, V., and Rombokas, E. (2018). “Evaluation of a visual localization system for environment awareness in assistive devices,” in Annual International Conference of the IEEE Engineering in Medicine and Biology Society (EMBC) (Honolulu, HI: IEEE), 5135–5141. doi: 10.1109/EMBC.2018.8513442.

Sandler, M., Howard, A., Zhu, M., Zhmoginov, A., and Chen, L. C. (2018). MobileNetV2: Inverted residuals and linear bottlenecks. arXiv [Preprint]. arXiv:1801.04381.

Simonyan, K., and Zisserman, A. (2014). Very deep convolutional networks for large-scale image recognition. arXiv [Preprint]. arXiv:1409.1556.

Spanias, J. A., Simon, A. M., Ingraham, K. A., and Hargrove, L. J. (2015). “Effect of additional mechanical sensor data on an EMG-based pattern recognition system for a powered leg prosthesis,” in International IEEE/EMBS Conference on Neural Engineering (NER) (Montpellier, IEEE), 639–642. doi: 10.1109/NER.2015.7146704.

Szegedy, C., Vanhoucke, V., Ioffe, S., Shlens, J., and Wojna, Z. (2015). Rethinking the Inception architecture for computer vision. arXiv [Preprint]. arXiv:1512.00567.

Tan, M., and Le, Q. V. (2019). EfficientNet: Rethinking model scaling for convolutional neural networks. arXiv [Preprint]. arXiv:1905.11946.

Tucker, M. R., Olivier, J., Pagel, A., Bleuler, H., Bouri, M., Lambercy, O., et al. (2015). Control strategies for active lower extremity prosthetics and orthotics: A review. Journal of NeuroEngineering and Rehabilitation, 12, 1. doi: 10.1186/1743-0003-12-1.

Varol, H. A., Sup, F., and Goldfarb, M. (2010). Multiclass real-time intent recognition of a powered lower limb prosthesis. IEEE Transactions on Biomedical Engineering, 57, 542–551. doi: 10.1109/TBME.2009.2034734.

Varol, H. A., and Massalin, Y. (2016). “A feasibility study of depth image based intent recognition for lower limb prostheses,” in Annual International Conference of the IEEE Engineering in Medicine and Biology Society (EMBC) (Orlando, FL: IEEE), 5055–5058. doi: 10.1109/EMBC.2016.7591863.

Vaswani, A., Shazeer, N., Parmar, N., Uszkoreit, J., Jones, L., Gomez, A. N., Kaiser, L., and Polosukhin, I. (2017). Attention is all you need. arXiv [Preprint]. arXiv:1706.03762.

Wang, D., Du, L., and Huang, H. (2013). “Terrain recognition improves the performance of neural-machine interface for locomotion mode recognition,” in IEEE International Conference on Computing, Networking and Communications (ICNC) (San Diego, CA: IEEE), 87–91. doi: 10.1109/ICCNC.2013.6504059.

Wong, A. (2018). NetScore: Towards universal metrics for large-scale performance analysis of deep neural networks for practical on-device edge usage. arXiv [Preprint]. arXiv:1806.05512.

Young, A. J., and Ferris, D. P. (2017). State of the art and future directions for lower limb robotic exoskeletons. IEEE Transactions on Neural Systems and Rehabilitation Engineering, 25, 171–182. doi: 10.1109/TNSRE.2016.2521160.

Zhang, F., Fang, Z., Liu, M., and Huang, H. (2011). “Preliminary design of a terrain recognition system,” in Annual International Conference of the IEEE Engineering in Medicine and Biology Society (EMBC) (Boston, MA: IEEE), 5452–5455. doi: 10.1109/IEMBS.2011.6091391.

Zhang, K., De Silva, C. W., and Fu, C. (2019a). Sensor fusion for predictive control of human-prosthesis-environment dynamics in assistive walking: A survey. arXiv [Preprint]. arXiv:1903.07674.

Zhang, K., Zhang, W., Xiao, W., Liu, H., De Silva, C. W., and Fu, C. (2019b). Sequential decision fusion for environmental classification in assistive walking. IEEE Transactions on Neural Systems and Rehabilitation Engineering, 27, 1780–1790. doi: 10.1109/TNSRE.2019.2935765.

Zhang, K., Xiong, C., Zhang, W., Liu, H., Lai, D., Rong, Y., et al. (2019c). Environmental features recognition for lower limb prostheses toward predictive walking. IEEE Transactions on Neural Systems and Rehabilitation Engineering, 27, 465–476. doi: 10.1109/TNSRE.2019.2895221.

Zhang, K., Wang, J., and Fu, C. (2019d). Directional PointNet: 3D environmental classification for wearable robotics. arXiv [Preprint]. arXiv: 1903.06846.

Zhang, K., Luo, J., Xiao, W., Zhang, W., Liu, H., Zhu, J., et al. (2020). A subvision system for enhancing the environmental adaptability of the powered transfemoral prosthesis. IEEE Transactions on Cybernetics. doi: 10.1109/TCYB.2020.2978216.

Zhong, B., Da Silva, R. L., Li, M., Huang, H., and Lobaton, E. (2020). Environmental context prediction for lower limb prostheses with uncertainty quantification. IEEE Transactions on Automation Science and Engineering. doi: 10.1109/TASE.2020.2993399.

Ziegler-Graham, K., MacKenzie, E. J., Ephraim, P. L., Travison, T. G., and Brookmeyer, R. (2008). Estimating the prevalence of limb loss in the United States: 2005 to 2050. Archives of Physical Medicine and Rehabilitation, 89, 422–429. doi: 10.1016/j.apmr.2007.11.005.

